# PIP4K2C inhibition reverses autophagic flux impairment induced by SARS-CoV-2

**DOI:** 10.1101/2024.04.15.589676

**Authors:** Marwah Karim, Manjari Mishra, Chieh-Wen Lo, Sirle Saul, Halise Busra Cagirici, Do Hoang Nhu Tran, Aditi Agrawal, Luca Ghita, Amrita Ojha, Michael P. East, Karen Anbro Gammeltoft, Malaya Kumar Sahoo, Gary L. Johnson, Soumita Das, Dirk Jochmans, Courtney A. Cohen, Judith Gottwein, John Dye, Norma Neff, Benjamin A. Pinsky, Tuomo Laitinen, Tatu Pantsar, Antti Poso, Fabio Zanini, Steven De Jonghe, Christopher R M Asquith, Shirit Einav

## Abstract

In search for broad-spectrum antivirals, we discovered a small molecule inhibitor, RMC-113, that potently suppresses the replication of multiple RNA viruses including SARS-CoV-2 in human lung organoids. We demonstrated selective dual inhibition of the lipid kinases PIP4K2C and PIKfyve by RMC-113 and target engagement by its clickable analog. Advanced lipidomics revealed alteration of SARS-CoV-2-induced phosphoinositide signature by RMC-113 and linked its antiviral effect with functional PIP4K2C and PIKfyve inhibition. We discovered PIP4K2C’s roles in SARS-CoV-2 entry, RNA replication, and assembly/egress, validating it as a druggable antiviral target. Integrating proteomics, single-cell transcriptomics, and functional assays revealed that PIP4K2C binds SARS-CoV-2 nonstructural protein 6 and regulates virus-induced impairment of autophagic flux. Reversing this autophagic flux impairment is a mechanism of antiviral action of RMC-113. These findings reveal virus-induced autophagy regulation via PIP4K2C, an understudied kinase, and propose dual inhibition of PIP4K2C and PIKfyve as a candidate strategy to combat emerging viruses.

## Introduction

Emerging viral infections pose major threats to human health. Severe acute respiratory syndrome coronavirus 2 (SARS-CoV-2) has resulted in over six million deaths. The incidence of mosquito-borne viral infections, such as those caused by the flavivirus dengue (DENV) and the alphavirus Venezuelan equine encephalitis virus (VEEV), has been increasing in part due to global warming^1^. The filoviruses Ebola (EBOV) and Marburg (MARV), the causative agents of lethal hemorrhagic fever, continue to cause sporadic outbreaks^2^. No effective countermeasures are currently available for the majority of these and other emerging viral infections. The current prevailing approved strategy—largely relying on targeting viral factors by direct-acting antivirals (DAAs)—is typically limited by a narrow spectrum of coverage and the emergence of drug resistance. There is thus a large unmet need for novel approaches, to be used individually or in combination with DAAs. Targeting cellular kinases exploited by multiple viruses is one attractive approach to overcome these challenges and provide readiness for future outbreaks, with demonstrated preclinical feasibility^3, 4^.

Phosphoinositides comprise a family of the membrane lipid phosphatidylinositol (PI) and its seven phosphorylation products, whose activity is tightly regulated by cellular kinases^5^. PI-3,5- bisphosphate [PI(3,5)P_2_] is generated through 5’-phosphorylation of PI-3-monophosphate [PI(3)P] by PIKfyve (PI-3-phosphate 5-kinase) on late endosomal/lysosomal and autophagic compartments^6^. Depletion of PI(3,5)P_2_ via PIKfyve inhibition disrupts lysosomal function, leading to their enlargement^7^. Yet, this effect is non-cytotoxic, as evidenced by the favorable safety profile demonstrated in clinical trials for inflammatory diseases with apilimod, a selective PIKfyve inhibitor^8–10^. PIKfyve is required for viral infections, and inhibiting it pharmacologically suppresses viral entry (coronaviruses, filoviruses) and/or egress (filoviruses, Lassa virus)^11–13^.

PI-4,5-bisphosphate [PI(4,5)P_2_] is formed via 4’-phosphorylation of PI5P by PI-5-phosphate 4-kinases: products of the *PIP4K2A, PIP4K2B*, and *PIP4K2C* genes^14^. These isoforms, sharing over 60% similarity, exhibit differences in enzymatic activity levels and subcellular distribution^15^. Whereas PIP4K2A/B positively regulates autophagic flux, PIP4K2C knockdown was shown to reduce the levels of an autophagy cargo protein and mutant huntingtin protein (mHTT) and to positively regulate mTORC1^15–19^ Yet, the biology of PIP4K2C is poorly understood, its role in viral infections remains unexplored, and its therapeutic relevance remains unknown.

Various viruses Impair autophagic flux to support their replication^20–23^. The SARS-CoV-2 nonstructural 6 protein (NSP6) was shown to bind the ATP6AP1 subunit of the vacuolar ATPase, disrupting lysosomal acidification and autophagic flux^24^. However, impaired autophagic flux is currently not being targeted pharmacologically. While optimizing inhibitors of the NUMB-associated kinase GAK as an antiviral strategy^25, 26^, we identified a novel small molecule inhibitor, RMC-113, that did not suppress GAK activity. Here, we discover that RMC-113 selectively inhibits PIP4K2C and PIKfyve and suppresses the replication of multiple RNA viruses. We probe the roles of these kinases in SARS-CoV-2 infection and the therapeutic potential and mechanism of antiviral action of RMC-113. Our findings validate PIP4K2C as a druggable antiviral target, beyond PIKfyve, demonstrating its binding to NSP6 and regulation at various stages of the SARS-CoV-2 life cycle, in-part by controlling viral-induced impairment of autophagic flux. Moreover, our findings highlight PIP4K2C and PIKfyve as the molecular targets of RMC-113 and enhancement of autophagic flux as its mechanism of antiviral action, thus proposing a candidate broad-spectrum antiviral for further development.

## Results

### RMC-113 demonstrates broad-spectrum antiviral activity in vitro and in human adult lung organoid (ALO)-derived monolayers with a high genetic barrier to resistance

Structure-affinity relationship analysis of isothiazolo[4,3*-b*]pyridine-based analogs for targeting GAK^25, 26^, generated a 3,6-disubstituted analog, RMC-113, which exhibited low affinity to GAK (K_d_=7.6 µM) (Fig. 1a). To determine the antiviral potential of RMC-113, we studied its effect against SARS-CoV-2. Five-day treatment with RMC-113 dose-dependently rescued Vero E6 cells constitutively expressing enhanced green fluorescent protein (eGFP) from SARS-CoV-2-induced lethality (isolate: Belgium-GHB-03021) (Fig. 1b-d). Similarly, RMC-113 dose-dependently inhibited replication of WT and SARS-CoV-2 expressing Nluc-reporter (rSARS-CoV-2/Nluc (USA-WA1/2020 strain)) in human lung epithelial (Calu-3) cells, as measured via plaque (EC_50_=0.25 μM) and luciferase (EC_50_=1.45 μM) assays, respectively (Fig. 1e and Supplementary Fig. 1a). It also suppressed replication-restricted pseudovirus bearing SARS-CoV-2 spike (S) protein (rVSV-SARS-CoV-2-S) in Vero cells (EC_50_=1.8 μM) (Supplementary Fig. 1b). No apparent effect on cellular viability was measured at the concentrations used in the infected cells via alamarBlue assays (CC_50_> 20 μM) (Fig. 1e and Supplementary Fig. 1a,b). Beyond SARS-CoV-2, RMC-113 dose-dependently inhibited the replication of the vaccine strain of VEEV (TC-83) via luciferase assays in human astrocytes (U-87 MG) (EC_50_=1.4 µM), DENV2 (EC_50_=1.4 µM) via plaque assays, and EBOV (EC_50_=5 μM) and MARV (EC_50_=7.8 μM) via microneutralization assays in human hepatoma (Huh7) cells, with no apparent cellular toxicity (Supplementary Fig. 1c-f).

**Fig. 1:**
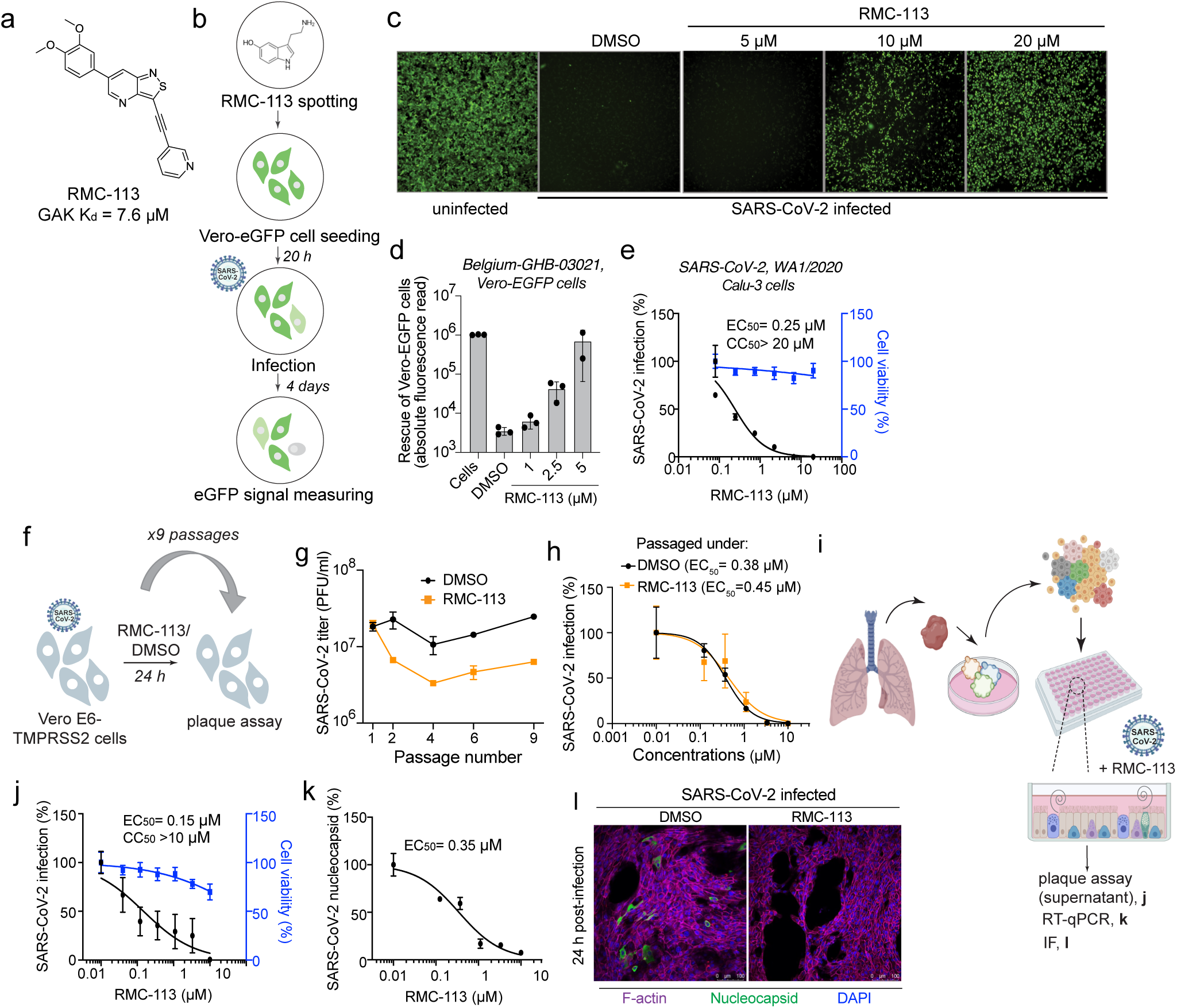
RMC-113 inhibits SARS-CoV-2 infection in vitro and in human ALOs with a high genetic barrier to resistance. **a,** Chemical structure of RMC-113. **b,** Rescue assay for virus-induced cell lethality. RMC-113 (10 μM) was incubated with Vero E6-eGFP cells for 20 hours followed by SARS-CoV-2 infection. eGFP signal measured at 96 hpi indicates cell survival. **c** and **d,** Fluorescence images (**c**) and corresponding graph (**d**) of Vero-eGFP cells rescued from SARS-CoV-2-induced lethality by RMC-113 (Belgium-GHB-03021 strain, MOI = 0.05). **e** and **j,** Dose response to RMC-113 of SARS-CoV-2 infection [black, USA-WA1/2020 strain, MOI = 0.05 (**e**), 1 (**j**)] and cell viability (blue) in Calu-3 cells (**e**) or ALO-derived monolayer supernatants (**j**) via plaque and alamarBlue assays at 24 (**e**) or 48 (**j**) hpi, respectively. **f** and **i,** Schematics of the experiments shown in **g** and **j, k, l**, respectively. **g,** Vero E6-TMPRSS2 cells were infected with rSARS-CoV-2-nLuc virus (MOI = 0.05) and passaged daily under RMC-113 (0.1-0.3 μM) or DMSO over nine passages. Viral titers were measured by plaque assays. **h,** Dose response to RMC-113 of rSARS-CoV-2-nLuc virus harvested after nine passages under RMC-113 or DMSO via luciferase assays. **k,** Dose response to RMC-113 of SARS-CoV-2 (MOI = 1) nucleocapsid copy number in ALO lysates measured by RT-qPCR assays at 48 hpi. **l,** Confocal IF microscopy images of F-actin (violet), nucleocapsid (green), and DAPI (blue) in naive and SARS-CoV-2–infected ALOs, pretreated with DMSO or RMC-113 (5 μM) at 24 hpi. Representative merged images at x40 magnification are shown. Scale bars: 50 μm. Data is representative (**c, d, g, j, l**), or a combination (**e, h, k**) of 2 independent experiments, each with 2–3 biological replicates. Data in **e, h, j,** and **k** are relative to DMSO. Means ± SD are shown.

To assess the barrier to resistance, SARS-CoV-2 was passaged in TMPRSS2 expressing Vero E6 (Vero E6-TMPRSS2) cells in the presence of RMC-113 at concentrations corresponding to values between the EC_50_ and EC_90_ or DMSO, and infectious viral titers were measured in culture supernatants by plaque assays (Fig. 1f). No phenotypic resistance was observed over nine passages (Fig. 1g). Furthermore, SARS-CoV-2 harvested following nine passages under RMC-113 treatment remained susceptible to RMC-113 (Fig. 1h).

Next, we studied the effect of RMC-113 treatment on SARS-CoV-2 infection in human adult stem cell-derived lung organoid (ALO)-monolayers composed of airway and alveolar cells^3, 27^ (Fig. 1i). RMC-113 dose-dependently suppressed SARS-CoV-2 replication measured in ALO culture supernatants by plaque assays (EC_50_=0.15 μM) and nucleocapsid (N) transcript expression measured in ALO lysates by RT-qPCR (EC_50_=0.35 μM), and CC_50_>10 μM (Fig. 1j,k). Moreover, RMC-113 treatment nearly abolished SARS-CoV-2 nucleocapsid (N) expression, as shown by confocal immunofluorescence (IF) analysis (Fig. 1l).

These results provide evidence that RMC-113 is a broad-spectrum antiviral candidate and suggest it targets a cellular function distinct from GAK.

### RMC-113 selectively inhibits PIP4K2C and PIKfyve

To identify the cellular targets, we first conducted kinome profiling in cell lysates treated with RMC-113 via multiplexed inhibitor beads kinome profiling coupled with mass spectrometry (MIB/MS)^28, 29^. RMC-113 exhibited dose-dependent binding of the lipid kinases PIKfyve, PIP4K2A, PIP4K2B, and PIP4K2C, with low or no affinity for other kinases and overall excellent selectivity (Fig. 2a and Supplementary Fig. 2a). A radiometric kinase activity screening of a 335-kinase panel that does not include PIKfyve and PIP4K2s measured no activity against any kinase, with an excellent selectivity score (S(<50%)=0.003) (Supplementary Fig. 2b,c and Table S1). RMC-113 potently bound recombinant PIKfyve (K_d_=370 nM) and PIP4K2C (K_d_=46 nM) proteins and suppressed the enzymatic activity of PIKfyve (IC_50_=8 nM) (Fig. 2b,c). While no in vitro enzymatic assay is currently available for PIP4K2C, cell-based target engagement analysis via live-cell NanoBRET assays revealed comparable activities on PIP4K2C and PIKfyve (IC_50_=392 nM and IC_50_=299.8 nM, respectively) (Fig. 2d)^30^. No kinase activity (IC_50_>10 µM) and lower affinity (K_d_=1.7 µM) were measured on PIP4K2A and PIP4K2B, respectively.

**Fig. 2:**
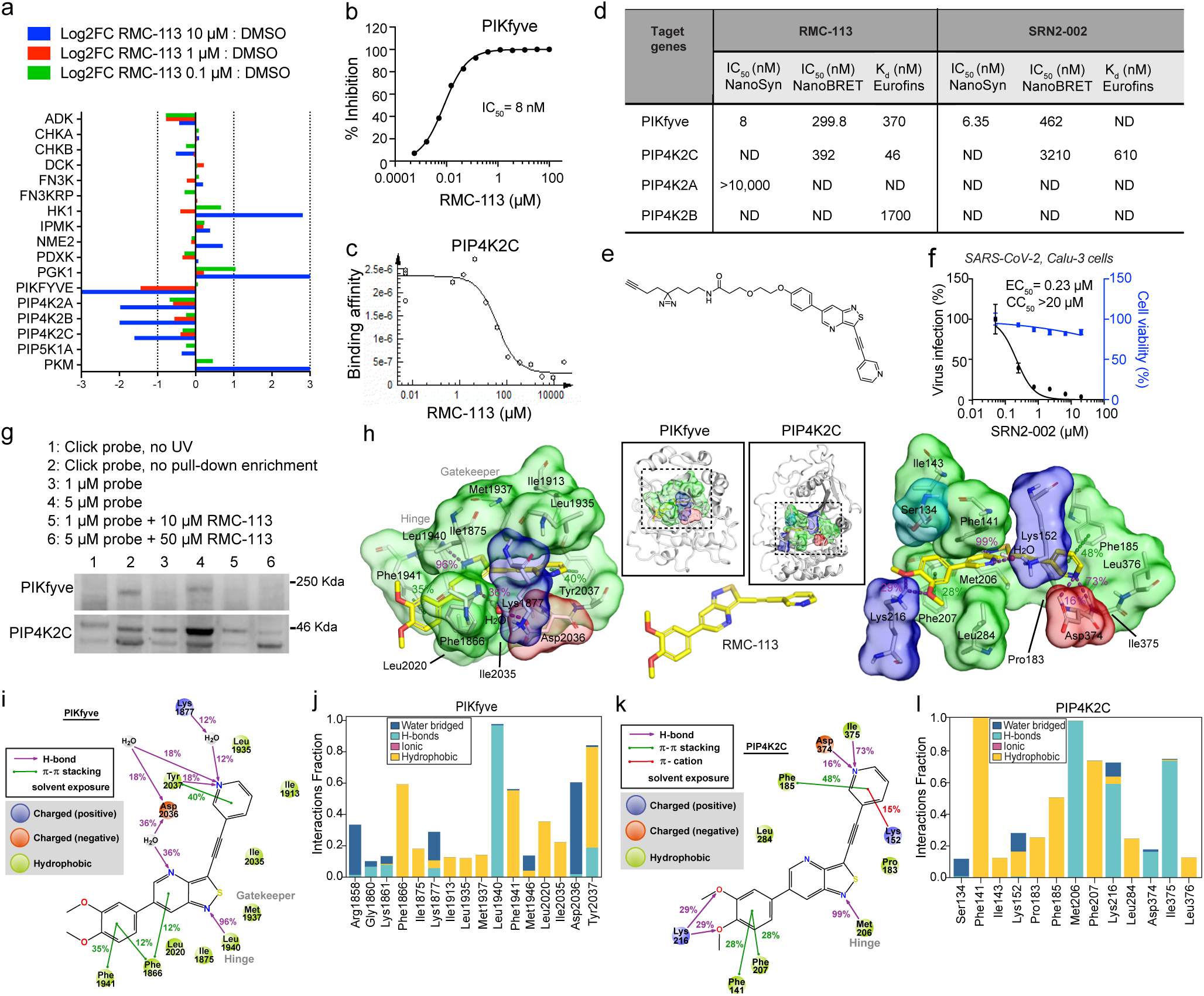
RMC-113 is a selective kinase inhibitor that stably binds PIKfyve and PIP4K2C. **a,** Kinase abundance ratio between RMC-113 (0.1, 1.0, and 10 µM)- and DMSO-treated SUM159 cell lysates measured following affinity purification via multiplexed inhibitor beads kinome profiling and analyzed by mass spectrometry (MIB/MS). Shown are the Log2 fold change values of a subset of the screen panel (see **Supplementary Fig. 2a**). Data means ± SD of 2 replicates are shown. **b** and **c,** In vitro dose response to RMC-113 of PIKfyve activity (**b**) and PIP4K2C binding (**c**). **d,** Biochemical parameters for RMC-113 and SRN2-002. ND=Not determined **e,** Chemical structure of SRN2-002. **f,** Dose response to SRN2-002 of SARS-CoV-2 (MOI = 0.05) infection (black) and cell viability (blue) in Calu-3 cells via plaque and alamarBlue assays at 24 hpi, respectively. **g,** PIKfyve and PIP4K2C expression in lysates of SARS-CoV-2-infected A549-ACE2 cells following incubation with SRN2-002 individually or in combination with RMC-113, UV irradiation, and pull down by streptavidin beads measured via Western blotting at 24 hpi. Shown are representative membranes of 2 experiments. Lanes: 1: SRN2-002, no UV; 2: SRN2-002, no pull-down enrichment; 3: SRN2-002 (1 µM); 4: SRN2-002 (5 µM); 5: competitive inhibition of SRN2-002 (1 µM) with RMC-113 (10 µM); 6: competitive inhibition of SRN2-002 (5 µM) with RMC-113 (50 µM). **h,** Putative binding mode of RMC-113 into the ATP-binding pockets of PIKfyve and PIP4K2C based on microsecond timescale MD simulations. A representative snapshot with key interactions are shown. Binding pocket residues with >10% interaction frequencies to RMC-113 are shown. Residues with positive charge are highlighted with blue molecular surface; negative charge, red; hydrophobic, green; polar, cyan. RMC-113 is shown in yellow (carbon) stick model. H-bonds are illustrated with purple dashed lines, ν–ν interactions with green. See **Supplemental Fig. 2f** for more detailed simulation interactions. **i** and **k**, Summary of the observed main protein-ligand interactions in the MD simulations of PIKfyve (i) and PIP4K2C (k). Interactions with >10% frequency are displayed. **j** and **l,** Aggregate of protein-ligand interactions (residues with >10%) in the simulations of PIKfyve (j) and PIP4K2C (l). Data in (**i** and **j)** consist of 72 μs, and (**k** and **l)** consist of 20 μs, both analyzed each 1 ns.

To validate these targets, we designed a clickable analogue, SRN2-002, by adding a terminal alkynyl motif with an aziridine photoaffinity group *via* an ethylene glycol linker, replacing the solvent-exposed 3,4-dimethoxyphenyl moiety^31^ (Fig. 2e). SRN2-002 exhibited potent activity on PIKfyve (in vitro: IC_50_=6.35 nM; cell-based: IC_50_=462 nM) and moderate activity on PIP4K2C activity (in vitro: K_d_=610 nM; cell-based: IC_50_=3210 nM) (Fig. 2d). SRN2-002 dose-dependently suppressed SARS-CoV-2 infection in Calu-3 cells with EC_50_ and CC_50_ values (EC_50_=0.23 µM, CC_50_>20 µM) comparable to those of RMC-113 (Fig. 2f). We confirmed the copper-catalyzed azide-alkyne cycloaddition CuAAC/click reaction between SRN2-002’s alkyne and azide-biotin (Supplementary Fig. 2d,e). Induced fit docking suggested that binding of SRN2-002 to the ATP-binding site of the kinases is plausible (Supplemental Fig. 2f,g) and that its reduced PIP4K2C activity may be due to the tighter environment with the solvent-oriented clickable group.

To assess target engagement, A549-ACE2 cells were infected with SARS-CoV-2 and treated with SRN2-002 alone or in combination with (non-clickable) RMC-113. After UV irradiation of cell lysates to bind the click probe to targets and a click reaction, pull-down assays using streptavidin beads were conducted followed by Western blot analysis. SRN2-002 at 5 µM pulled down PIKfyve and PIP4K2C, with minimal signal in non-UV irradiated samples. Addition of RMC-113 dose-dependently reduced PIKfyve and PIP4K2C pull-down, indicating effective competition and target engagement (Fig. 2g).

Collectively, these findings provide evidence that RMC-113 is a dual inhibitor of PIP4K2C and PIKfyve.

### RMC-113 displays comparable binding at the active site of PIP4K2C and PIKfyve

To disclose the putative binding mode of RMC-113 to these kinases, we utilized a molecular dynamics (MD) simulation approach^32^. Our microsecond timescale simulations suggest a plausible reasonable binding to RMC-113 in the ATP-binding site of both PIKfyve and PIP4K2C (Fig. 2h). Both kinases had a comparable binding mode with RMC-113 based on their simulation interactions, with a constant hydrogen-bond to the hinge region throughout the simulations (PIP4K2C: Met206; PIKfyve: Leu1940). Two phenylalanine residues located above and below accommodate the aromatic dimethoxyphenyl in the binding pocket (PIP4K2C: Phe141, Phe207; PIKfyve: Phe1866, Phe1941). On the pyridine-end of the molecule, p–p stacking is observed with an aromatic residue (PIP4K2C: Phe185; PIKfyve: Tyr2037). A key difference in the interactions of the solvent-exposed region between the kinases is that whereas interactions with Lys216 are observed in PIP4K2C, no stable interactions are observed in this location in PIKfyve. The behavior of RMC-113 in complex with these two kinases was broadly similar, with the one key movement within the binding being a flip with the solvent-exposed dimethoxyphenyl-groups (Fig. 2i,j and Supplementary Fig. 3a,b). The pyridine-end of the molecule appears slightly more stable with PIP4KC, most likely due to the direct hydrogen-bond interactions to its nitrogen (Fig. 2k,l and Supplementary Fig. 3a,b). Notably, both kinase domains display high flexibility in the simulations, highlighting their specific characteristics (Supplementary Fig. 3a-d).

Overall, these simulations imply a stable binding of RMC-113 to both kinases, consistent with the experimental data.

### PIKfyve and PIP4K2C are required for effective SARS-CoV-2 infection and are the molecular targets mediating the antiviral effect of RMC-113

To define the molecular targets mediating RMC-113’s antiviral effect, we assessed the effect of siRNAs targeting PIKfyve and PIP4K2C on SARS-CoV-2 in Calu-3 cells via plaque assays. Knockdown efficiency was confirmed via RT-qPCR (Fig. 3a,b), and alamarBlue assays showed no cellular toxicity (Fig. 3c). Depletion of PIKfyve and PIP4K2C suppressed SARS-CoV-2 replication by over 2 logs relative to a non-targeting (siNT) control (Fig. 3c).

**Fig. 3:**
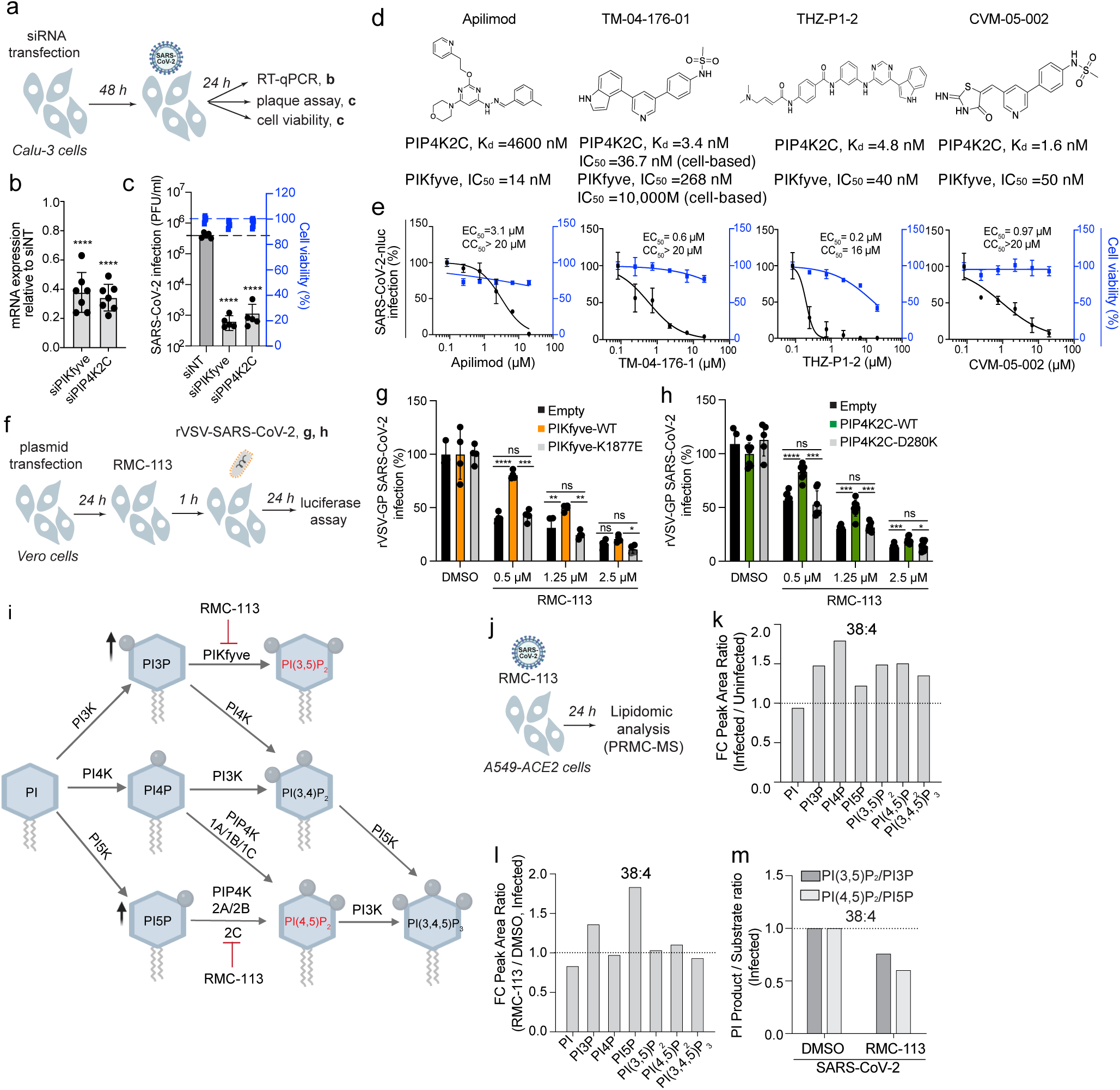
PIKfyve and PIP4K2C are essential for SARS-CoV-2 infection and mediate the antiviral effect of RMC-113. a, f, j. Schematics of the experiments in **b**, **c** (**a**); **g**, **h** (**f**); and **k, l, m** (**j**). **b,** Confirmation of siRNA-mediated knockdown by RT-qPCR in Calu-3 cells. Shown are gene expression levels normalized to GAPDH relative to respective gene levels in siNT control at 48 hours post-transfection. **c,** Viral titers (PFU/ml) and cell viability (blue) in Calu-3 cells transfected with the indicated siRNAs at 24 hpi with SARS-CoV-2 (MOI = 0.05) via plaque and alamarBlue assays, respectively. **d,** Structures of PIKfyve and/or PIP4K2C inhibitors. **e,** Dose response of rSARS-CoV-2-nLuc (black, USA-WA1/2020 strain, MOI = 0.05) infection and cell viability in Calu-3 cells via luciferase and alamarBlue assays at 24 hpi, respectively. **g** and **h,** Rescue of rVSV-SARSCoV-2-S infection under RMC-113 treatment upon ectopic expression of WT PIKfyve (**g**) and PIP4K2C (**h**), their kinase-dead mutants or empty control plasmids measured by luciferase assays at 24 hpi in Vero cells. **i**, Phosphoinositides and associated kinases. **k** and **l,** Fold change (FC) of peak area ratio of the indicated phosphoinositides in SARS-CoV-2-infected (MOI = 0.5) vs. uninfected (**k**) and RMC-113- vs. DMSO-treated infected Calu-3 cells (**l**), as measured via PRMC-MS. **m**, Product-to-substrate ratios in RMC-113- vs. DMSO-treated cells infected with SARS-CoV-2 (**k, l**). Data is relative to siNT (**b, c**) or DMSO (**e, g, h**). Data are a combination (**b**, **c**, **e**, **g,** and **h**) or representative of 2 independent experiments with 2-4 replicates each. Means ± SD are shown (**b, c, g, h**). **k, l, m** represent one of two independent experiments. See an associated experiment in **Supplemental Fig. 5e-g** and **Table S2.** *P ≤ 0.05, **P < 0.01, ***P <0.001, ****P <0.0001 by 1-way ANOVA followed by Dunnett’s (**b, c**) or Tukey’s multiple-comparison test (**g, h**).

We assessed the requirement of these kinases by testing the anti-SARS-CoV-2 effects of chemically distinct investigational compounds targeting PIKfyve and/or PIP4K2C (Fig. 3d). Apilimod, a selective PIKfyve inhibitor (In vitro: IC_50_=14 nM)^6^, dose-dependently inhibited the replication of SARS-CoV-2 in both Calu-3 cells (EC_50_=3.1 µM, CC_50_>20 µM), as reported^12^, and in ALOs (EC_50_=0.42 µM, CC_50_=9.9 µM)(Fig 3e and Supplementary Fig. 4a-c). Notably, TM-04-176-01, a selective PIP4K2C inhibitor (K_d_=3.4 nM; cell-based PIP4K2C: IC_50_=36.7 nM, PIKfyve: IC_50_=10,000 nM)^33^, potently suppressed SARS-CoV-2 replication (EC_50_=0.6 µM, CC_50_>20 µM) (Fig. 3e). THZ-P1-2 and CVM-05-002, inhibitors of PIKfyve and PIP4K2C^33, 34^, also dose-dependently inhibited SARS-CoV-2 infection (EC_50_=0.2 µM and EC_50_=0.97 µM, respectively), albeit THZ-P1-2 demonstrated greater cytotoxicity (CC_50_=16 µM) (Fig. 3e).

To verify that RMC-113’s antiviral mechanism is mediated by inhibition of the kinase activity of PIKfyve and PIP4K2C, we conducted “rescue” experiments in Vero cells infected with rVSV-SARS-CoV-2-S. Ectopic expression of wild-type (WT) PIKfyve or PIP4K2C (Supplementary Fig. 4d,e) completely or partially reversed the antiviral effect of RMC-113 (Fig. 3f-h and Supplemental Fig. 4f,g). Contrastingly, ectopic expression of catalytically inactive PIKfyve (K1877E) and PIP4K2C (D280K) mutants or a control plasmid did not reverse RMC-113’s antiviral effect (Fig. 3f-h and Supplemental Fig. 4f,g).

IF confocal microscopy analysis of A549-ACE2 cells ectopically expressing 2xFYVE, a PI3P marker fused to mCherry revealed induction of cytosolic vacuolation and swelling of membrane structures that stained positive for the endosomal markers EEA-1 and Rab-7 upon RMC-113 and apilimod treatment (Supplementary Fig. 4h-j), further supporting PIKfyve suppression with no apparent change in its subcellular distribution.

Unlike PIKfyve^12^, the functional relevance of PIP4K2C in viral infections has not been previously demonstrated. These findings provide genetic and pharmacological validation that PIP4K2C, beyond PIKfyve, is a druggable antiviral target and a molecular target mediating the antiviral effects of RMC-113.

### RMC-113 alters the phosphoinositide regioisomer signature by advanced lipidomics analysis

To determine whether RMC-113 treatment impacts phosphoinositide abundance, lipid extracts derived from uninfected and SARS-CoV-2-infected A549-ACE2 cells were subject to lipidomic analysis. Employing Phosphoinositide Regioisomer Measurement by Chiral column chromatography and Mass Spectrometry (PRMC-MS)^35^, we comprehensively profiled all eight PI classes and their acyl chain variants (defined by the carbon number and the saturation level) (Fig. 3i, j). Seven of the eight phosphoinositide classes (except for PI(3,4)P_2_) were detected in all tested conditions. Upon SARS-CoV-2 infection, the abundance of multiple PI classes was increased relative to uninfected samples, albeit with some variability across independent experiments (Fig. 3k and Supplementary Fig. 5a). This increase was most pronounced with the abundant acyl chain (38:4) (Fig. 3k), yet a similar trend was observed with other acyl chains (Supplementary Fig. 5b-e and Table S2). Notably, RMC-113 treatment in infected cells caused a 1.5-2-fold increase in the abundance of PI3P and PI5P—the substrates of PIKfyve and PIP4K2C, respectively^6, 36^—relative to DMSO controls (Fig. 3l and Supplementary Fig. 5f). No concomitant reduction in the levels of the respective phosphorylated products, PI(3,5)P_2_ and PI(4,5)P_2_, was detected in RMC-113- vs. DMSO-treated infected cells (Fig. 3l), likely due to intact activity of enzymes not targeted by RMC-113, such as PIP4K (1A/1B/1C)^37^ (Fig. 3i). Nevertheless, the product-to-substrate ratios of PIKfyve and PIP4K2C were reduced in both uninfected and infected cells upon RMC-113 treatment relative to DMSO (Fig. 3m and Supplementary Fig. 5g). Similar results were observed with other acyl chain variants (Table S2).

These findings provide evidence that the antiviral effect of RMC-113 is correlated with functional inhibition of PIP4K2C and PIKfyve activities and propose modulation of a virus-induced PI signature as a candidate mechanism of antiviral action.

### PIP4K2C is required for SARS-CoV-2 entry, RNA replication and assembly/egress, whereas PIKfyve is required for viral entry only

To pinpoint the steps of the viral life cycle impacted by RMC-113, we conducted time-of-addition experiments. RMC-113 was added to Calu-3 cells upon infection or at 2, 5, or 8 hpi with SARS-CoV-2 (Fig. 4a). Cell culture supernatants were harvested at 10 hpi, which represents a single cycle of viral replication in Calu-3 cells, and infectious viral titers were measured by plaque assays. RMC-113 treatment initiated upon infection onset and maintained throughout the 10-hour experiment (0 to 10) suppressed viral infection by 99.5% relative to DMSO control (Fig. 4b). RMC-113 treatment during the initial 2 hours of infection (0 to 2) suppressed viral infection by 79.4%, confirming an effect on entry of WT SARS-CoV-2 (beyond rVSV-SARS-CoV-2-S) (Fig. 4b and Supplementary Fig. 1b). Following extensive washing at 2 hpi (to remove the viral inoculum), the addition of RMC-113 at 2, 5, and 8 hpi suppressed viral infection by 97%, 96%, and 69.3%, respectively, indicating inhibition at post-entry stages (Fig. 4b). In contrast, treatment with apilimod suppressed SARS-CoV-2 replication when added during the first two hpi, but not at later time points (Fig. 4c).

**Fig. 4:**
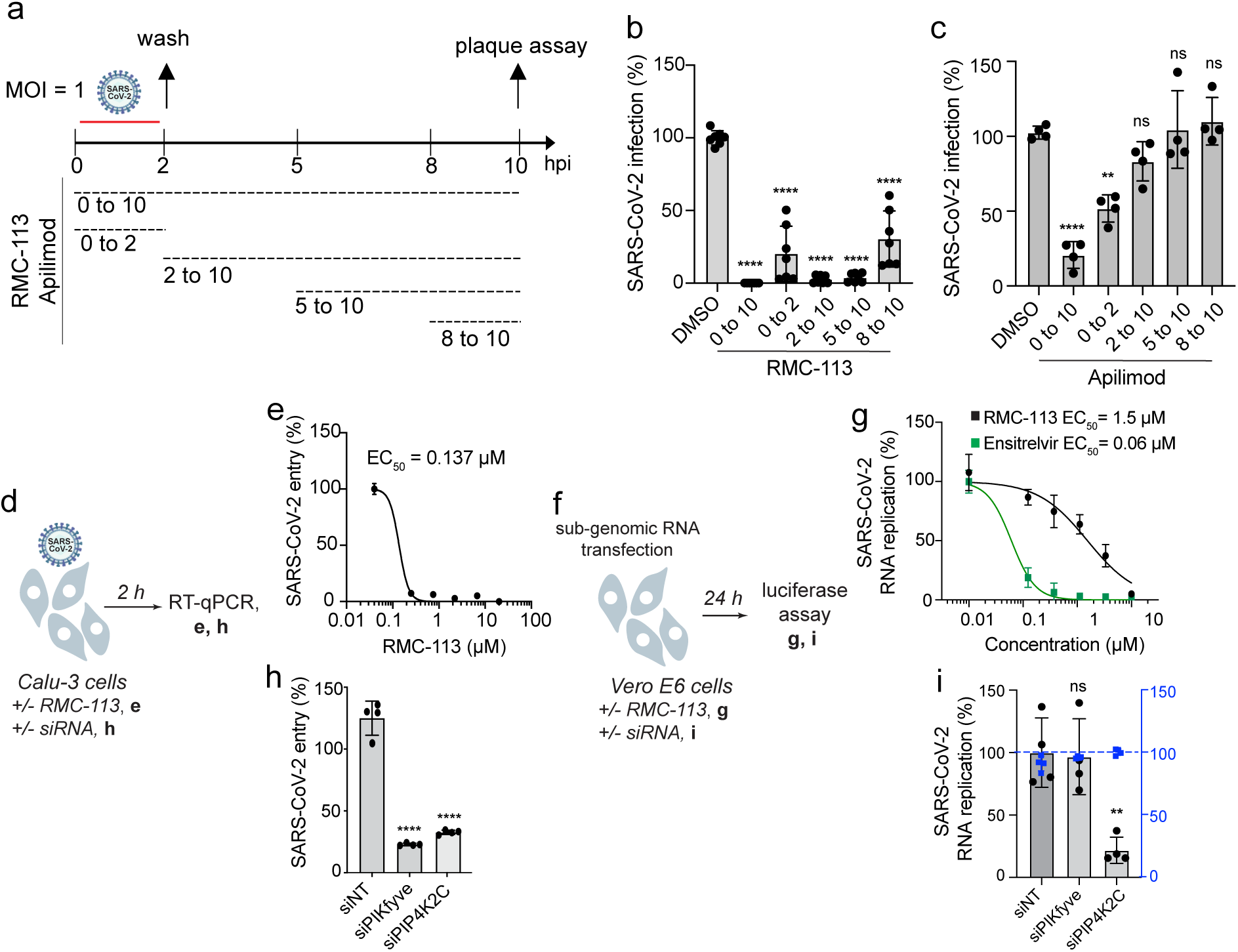
PIP4K2C is required for temporally distinct stages of the SARS-CoV-2 life cycle, whereas PIKfyve is required for viral entry only. **a,** Schematic of the time-of-addition experiments shown in **b**, **c**. **b** and **c,** Calu-3 cells were infected with WT SARS-CoV-2 (MOI = 1). At the indicated times, RMC-113 (10 μM) (**b**), apilimod (10 μM) (**c**), or DMSO were added. Supernatants were collected at 10 hpi, and viral titers measured by plaque assays. **d** and **f,** Schematics of the experiments shown in **e**, **h** (**d**) and **g, i** (**f**). **e,** Dose response to RMC-113 of WT SARS-CoV-2 entry (MOI = 1) in Calu-3 cell lysates measured by RT-qPCR assays at 2 hpi. **g,** Dose response to RMC-113 and ensitrelvir of viral RNA replication measured by luciferase assay in Vero E6 cells 24 hours post-transfection of in vitro transcribed nano-luciferase reporter-based SARS-CoV-2 subgenomic non-infectious replicon^38^. **h,** WT SARS-CoV-2 (MOI = 1) entry measured in Calu-3 cells depleted of the indicated kinases using corresponding siRNAs by RT-qPCR at 2 hpi. **i,** Viral RNA replication and cell viability (blue) measured by luciferase and alamarBlue assays, respectively, in Vero E6 cells depleted of the indicated kinases, 24 hours post-transfection of in vitro transcribed nano-luciferase reporter-based SARS-CoV-2 subgenomic non-infectious replicon^38^. Data are a combination (**b**, **c**, **e**, **g, h**) or representative (**i**) of two independent experiments with 2-4 replicates each. Means ± SD are shown. Data is relative to DMSO (**b, c, e, g**) or siNT (**h, i**). **P < 0.01, ****P <0.0001 by 1-way ANOVA followed by Turkey’s (**b,c**) or Dunnett’s (**h, i**) multiple-comparison test. ns= non-significant.

RMC-113 dose-dependently suppressed intracellular SARS-CoV-2 N copy number (EC_50_=0.137 μM) at 2 hpi of Calu-3 cells with a high-inoculum virus relative to DMSO, as measured by RT-qPCR, validating an effect on SARS-CoV-2 entry (Fig. 4d,e). Moreover, RMC-113 treatment dose-dependently inhibited the replication of in vitro transcribed RNA encoding a nano-luciferase reporter-based SARS-CoV-2 subgenomic replicon deleted for the spike (S), envelope (E), and membrane (M) proteins (ΔS-E-M)^38^ in Vero E6 cells. Similar to ensitrelvir, albeit with a higher EC_50_ value (1.5 µM vs. 0.06 µM), this effect reveals that RMC-113 also suppresses viral RNA replication (Fig. 4f,g). In contrast, apilimod treatment suppressed rVSV-SARS-CoV-2-S pseudovirus infection (Supplementary Fig. 6a,b), confirming its effect on SARS-CoV-2 entry, yet it dose-dependently increased viral RNA replication (Supplementary Fig. 6c,d).

To probe the requirement for PIKfyve and PIP4K2C in viral entry and RNA replication, we conducted the assays described above in cells depleted for the individual lipid kinases by siRNAs (Fig. 4f). Depletion of PIKfyve and PIP4K2C suppressed SARS-CoV-2 entry by 81.7% and 73.8%, respectively, relative to siNT as measured at 2 hpi by RT-qPCR (Fig. 4d,h). Moreover, depletion of PIP4K2C, but not PIKfyve, suppressed replication of the subgenomic replicon (Fig. 4f,i).

These findings pinpoint the role of PIKfyve to SARS-CoV-2 entry only and discover a role for PIP4K2C in viral entry, RNA replication, and assembly/egress. RMC-113, but not apilimod, thus suppresses temporally distinct stages of the SARS-CoV-2 life cycle.

### RMC-113 reverses SARS-CoV-2-induced impairment of autophagic flux

To test the hypothesis that by suppressing PIP4K2C, RMC-113 modulates autophagy, we studied the effect of RMC-113 on autophagosome formation and autophagic flux via IF analysis of multiple single A549-ACE2 cells expressing the premo-GFP-RFP-LC3 reporter and stained for SARS-CoV-2 N (Fig. 5a). Since GFP fluorescence is quenched in the acidic lysosomes whereas RFP signal is stable, in this assay, autophagosomes appear as yellow (RFP+/GFP+) and autolysosomes as red (RFP+/GFP-) puncta (Fig. 5b, and Supplementary Fig. 7a). Upon SARS-CoV-2 infection, we measured a 2.4-fold increase in the number of yellow puncta and a 1.2-fold decrease in the number of red puncta relative to uninfected controls (Fig. 5c and Supplementary Fig. 7b). The autophagic flux (autolysosome-to-autophagosome or red-to-yellow puncta ratio) was 2.4 folds lower in infected vs. uninfected cells (Fig. 5c,d). Thus, SARS-CoV-2 infection impairs autophagic flux but not autophagosome formation.

**Fig. 5:**
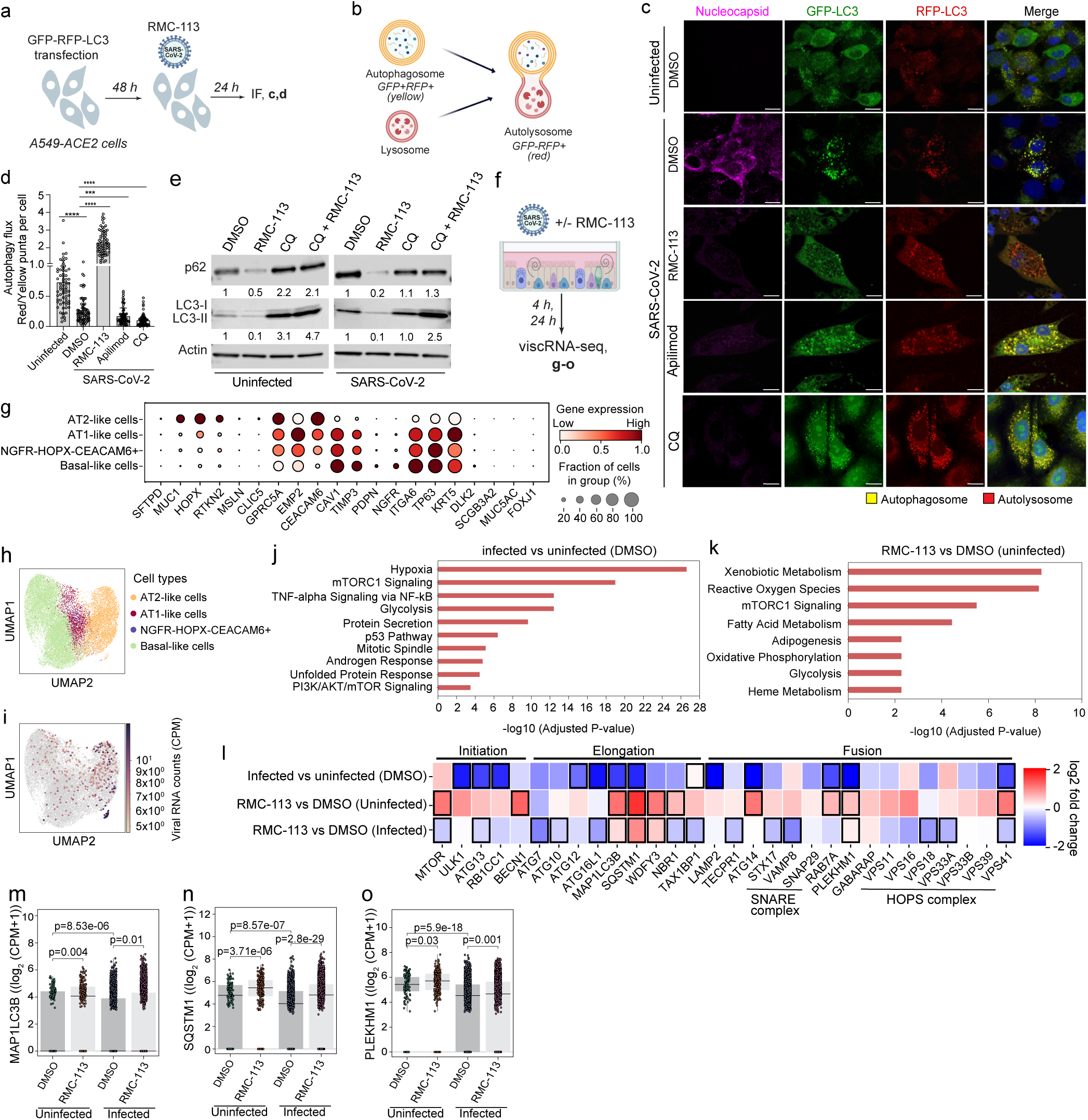
RMC-113 reverses SARS-CoV-2-induced impairment of autophagic flux. **a,** Schematic of the experiment shown in **c, d**. **b,** Autophagy flux: a shift from autophagosomes (yellow, RFP+/GFP+) to autolysosomes (red, RFP+/GFP-). **c,** Representative confocal microscopic images of A549-ACE2 cells transfected with GFP-RFP- LC3 tandem plasmid and infected with SARS-CoV-2 (MOI = 0.5), treated with DMSO, RMC-113 (5 μM), apilimod (5 μM) or CQ (1 μM) for 24 hours and stained for nucleocapsid (violet). Representative merged images at x40 magnification are shown Scale bars: 10 μm. Zoomed-in images show autophagosomes (yellow) and autolysosomes (red). **d,** Autolysosome-to-autophagosome ratio (autophagy flux) in 70 single cells (**c**). **e,** Expression levels of autophagy markers in uninfected and SARS-CoV-2-infected (MOI= 1) A549-ACE2 cell lysates treated with RMC-113 (5µM) and/or CQ (1µM) or DMSO at 24 hpi via Western blotting. Numbers represent expression signals relative to DMSO averaged from four membranes. **f,** Schematic of the viscRNA-seq analysis. **g,** Dot plot depicting marker genes used to annotate the indicated cell populations. Color indicates expression level in cpm; dot size indicates the fraction of cells expressing the marker. **h** and **i,** UMAP embedding of the scRNA-seq dataset indicating distinct cell types (**h**) or SARS-CoV-2 transcripts (**i**). **j** and **k,** Pathway enrichment in AT2 cells infected vs. uninfected (**j**) and RMC-113 vs. DMSO treated (**k**) ALOs. **l,** Heatmap showing the log2 fold change in the expression of autophagy-related genes between infected vs. uninfected (DMSO), RMC-113 vs. DMSO (uninfected); and RMC-113 vs. DMSO (infected) in AT2-like cells at 24 hpi. See, **Supplementary Fig. 8b**. Black rectangles highlight transcripts with significant changes measured by Wilcoxon test. **m-o,** Box plots showing the expression level of the indicated genes in individual AT2 cells at 24 hpi. Horizontal lines indicate the first, second (median) and third quartiles; whiskers extend to ±1.5× the interquartile range. P values by two-sided Wilcoxon test with Benjamini-Hochberg correction are shown. Data is a combination (**d, j-o,**) or representative (**c, e**) of 3 independent experiments. Means ± SD are shown (**d**). ***P < 0.001, ****P <0.0001 by 1-way ANOVA followed by Dunnett’s multiple comparison test (**d**).

RMC-113 treatment caused a 2-fold reduction in the number of yellow puncta, over 5-fold increase in the number of red puncta relative to DMSO (Supplementary Fig. 7b) and 8.8-fold increase in the autolysosome-to-autophagosome ratio, confirming that RMC-113 promotes autophagic flux (Fig. 5c,d). In contrast, apilimod and CQ did not alter autophagosome formation (Fig 5c and Supplementary Fig. 7b), but caused a 1.9- and 2.9-fold decrease in autolysosome numbers relative to DMSO, and a 1.6- and 2.8-fold reduction in the autolysosome-to-autophagosome ratio, respectively (Fig. 5c,d). Moreover, the colocalization of LC-3 with LAMP-1 (lysosomal marker) was greater in GFP-RFP-LC3-expressing cells upon treatment with RMC-113 than apilimod and CQ, suggesting increased autophagosome-lysosome fusion by this compound (Supplementary Fig. 7c).

Next, we assessed protein levels of p62 (autophagy cargo) and LC3-II (autophagic marker) in A549-ACE-2 cell lysates via Western blot analysis. The expression signals of p62 and LC3-II were greater in SARS-CoV-2-infected vs. uninfected (DMSO-treated) cells (Fig. 5e) suggesting that SARS-CoV-2 inhibits autophagic degradation, in agreement with prior studies^24, 39^. RMC-113 treatment caused a 2-10-fold decrease in the expression signals of p62 and LC3-II relative to DMSO in SARS-CoV-2-infected and uninfected cells (Fig. 5e), consistent with enhanced autophagic degradation. We also studied the effect of RMC-113 treatment independent of autophagosome turnover following treatment with chloroquine (CQ) an inhibitor of autophagosome-lysosome fusion^40^. While CQ did not significantly alter p62 and LC3-II expression signal in infected cells, the addition of RMC-113 following CQ treatment increased the LC3-II signal by 2.5 folds relative to DMSO (Fig. 5e), favoring increased autophagosome turnover upon RMC-113 treatment over reduced autophagosome formation^41^.

These findings provide evidence that RMC-113, but not apilimod, promotes autophagic degradation arrested by SARS-CoV-2 and this effect is mediated by PIP4K2C (vs. PIKfyve) inhibition.

### Virus-inclusive scRNA-seq (viscRNA-seq) analysis reveals temporal SARS-CoV-2-induced autophagy signatures and partial reversion by RMC-113

To determine whether the observed induction of autophagic degradation by RMC-113 is accompanied by transcriptional regulation of autophagy and/or lysosomal functions, we characterized the transcriptional host response to SARS-CoV-2 infection and RMC-113 treatment in correlation with viral RNA (vRNA) abundance via viscRNA-seq analysis^42^ using PARSE technology in ALOs at 4 and 24 hpi and treatment (Fig. 5f). We recovered 20,672 high-quality cells (Supplementary Fig. 8a) and identified four major cell populations: alveolar epithelial type II (AT2)-like, AT1-like, and basal-like cells, as well as NGFR-HOPX-CEACAM6+ cells, likely representing various differentiation stages (Fig. 5g,h). AT2-like cells were the main (poly-adenylated) viral RNA-harboring cells (VHCs) (Fig. 5i).

Gene ontology (GO) analysis of differentially expressed genes (DEGs) in AT2-like cells revealed mTORC1 signaling, an autophagy regulator^15, 18^, among the three top-upregulated pathways upon infection and RMC-113 treatment (Fig. 5j,k). We thus focused the differential expression analysis on genes involved in autophagy and lysosomal functions. At 4 hpi, AT2-like cells from SARS-CoV-2-infeced ALOs exhibited downregulation of *mTOR* along with upregulation of autophagy initiation genes *ULK1, ATG13, RB1CC1,* and *BECN1* relative to AT2-like cells from uninfected ALOs, and variable regulation of genes involved in elongation and fusion, suggesting infection-induced autophagy initiation. While a variable pattern was observed in uninfected ALOs, RMC-113 treatment of SARS-CoV-2-infected ALOs was associated with upregulation of *mTOR*, downregulation of *BECN1*, upregulation of elongation factors (e.g. *SQSTM1, MAP1LC3B*), and variable regulation of fusion genes, relative to DMSO controls (Supplementary Fig. 8b).

At 24 hpi, AT2-like cells from SARS-CoV-2-infected ALOs demonstrated mild upregulation of *mTOR* along with profound downregulation of multiple genes involved in autophagy initiation (*ULK1, ATG13, RB1CC1*), elongation [*ATG12, ATG16L1, MAP1LC3B (encoding p62), SQSTM1 (encoding LC3-II), TAX1BP1*] and fusion (*LAMP2, ATG14, RAB7A, PLEKHM1, VPS41*) relative to uninfected ALOs. In uninfected ALOs, RMC-113 treatment induced upregulation of genes involved in all stages of autophagy, particularly elongation (*MAP1LC3B, SQSTM1, WDFY3, NBR1*) and fusion [*ATG14, RAB7A, PLEKHM1* and member of the SNARE (*VAMP8*) and HOPS (*VPS41*) complexes] relative to DMSO (Fig. 5l). In SARS-CoV-2-infected ALOs, RMC-113 treatment resulted in a more variable signature with upregulation of some elongation (*MAP1LC3B, SQSTM1, and WDFY3*) and fusion (*PLEKHM1, GRABARAP)* genes, yet downregulation of others, as the SNARE (*STX17, VAMP8*) and HOPS (*VPS18, VSP33A* and *VSP41*) complex genes, relative to DMSO (Fig. 5l). Several genes including *MAP1LC3B*, *SQSTM1* and *PLEKHM1* demonstrated a pattern wherein expression level was significantly downregulated upon SARS-CoV-2 infection (vs. no infection) in DMSO-treated ALOs and significantly upregulated in response to RMC-113 treatment (Fig. 5m-o, and Supplementary Fig. 8c,d). The overall expression levels of autophagy genes was slightly higher in VHCs than bystander cells in both DMSO and RMC-113-treated ALOs (Supplementary Fig. 8e,f).

Downregulation of multiple genes involved in lysosomal functions was observed at 24 hpi in infected vs uninfected ALOs. These included membrane transporters, hydrolyses, multiple subunits of the v-ATPase complex and genes involved in lysosomal biogenesis and other functions (Supplementary Fig. 9). In contrast, RMC-113 treatment induced upregulation of multiple genes in the same categories, including membrane transporters (e.g. LAPTM4A, LMBRD1, LAMP1); hydrolyses (e.g. CTSZ, GGH, GNS), and v-ATPase complex subunits (Supplementary Fig. 9).

These findings indicate a transcriptional effect of SARS-CoV-2 infection on autophagocytic and lysosomal genes and proposes partial reversion of this phenotype as a potential mechanism of the antiviral action of RMC-113.

### PIP4K2C binds SARS-CoV-2 NSP6 and regulates SARS-CoV-2-induced impairment of autophagic flux

Repeating the functional autophagic flux experiments using a genetic approach revealed that PIP4K2C depletion caused a 41.6% reduction in the number of yellow puncta along with a 31.4% increase in the number of red puncta and a 2.6-fold increase in the autolysosome-to-autophagosome ratio relative to siNT control via IF analysis (Fig. 6a-c and Supplementary Fig. 10a-c). A similar, pattern was observed in uninfected cells (Supplementary Fig. 10b,c).

**Fig. 6:**
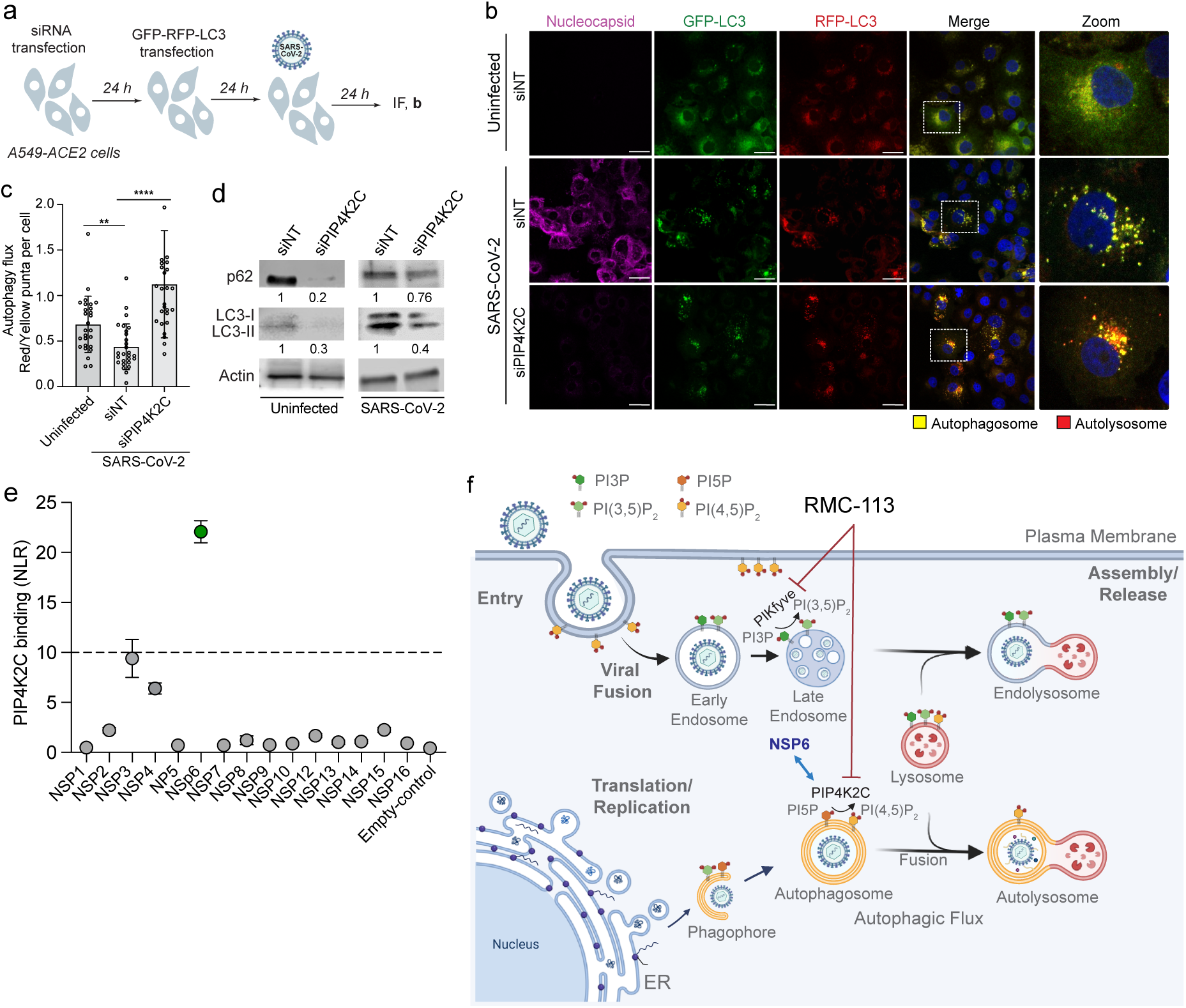
PIP4K2C binds SARS-CoV-2 NSP6 and mediates virus-induced impairment of autophagic flux. **a,** Schematic of the experiment shown in **b**. **b,** Representative confocal microscopic images of A549-ACE2 cells transfected with the indicated siRNAs and GFP-RFP-LC3 tandem plasmid, and infected with SARS-CoV-2 (MOI = 0.5) for 24 hours and stained for nucleocapsid (violet). Representative merged images at x40 magnification are shown. Scale bars: 10 μm. Zoomed-in images show autophagosomes (yellow) and autolysosomes (red). **c,** Autolysosome-to-autophagosomes ratio (autophagy flux) in 27 single cells (**b**). **d,** Expression levels of p62, LC3I and LC3II following transfection of the indicated siRNAs in uninfected A549-ACE2 cell lysates (left panel) and SARS-CoV-2-infected cells at 24 hpi (MOI = 0.5) (right panel). Numbers represent the expression signal relative to DMSO averaged from 3 membranes. **e,** PIP4K2C interactions with 15 SARS-CoV-2 nonstructural proteins and empty plasmid as measured via protein-fragment complementation assay (PCAs) in HEK 293T cells. Dots depict the mean normalized luminescence ratio (NLR) values generated from 2 independent experiments with 3 replicates each. The dotted line depicts the cutoff (NLR>10) used to define PIP4K2C-interacting proteins (depicted in green), representing greater than two SDs above the mean NLR of a non-interacting reference set. NLR, normalized luminescence ratio. **f,** Proposed model for the roles of PIP4K2C and PIKfyve in SARS-CoV-2 infection and the mechanism of antiviral action of RMC-113. Data is a representative (**b, d, e**) of 2 independent experiments. Data are relative to siNT (**c, d**). Means ± SD are shown (**c**). **P < 0.001, ****P <0.0001 by 1-way ANOVA followed by Dunnett’s multiple comparison test (**c**).

Concurrently, siRNA-mediated PIP4K2C depletion reduced the signals of p62 and LC3-II expression by 80% and 70% in uninfected A549-ACE2 cells and by 24% and 60% in SARS-CoV-2-infected cells, respectively, relative to siNT control as measured by Western blot analysis (Fig. 6d).

Lastly, to determine if PIP4K2C plays a direct role in SARS-CoV-2 infection, we screened for its interactions with 15 nonstructural SARS-CoV-2 proteins via protein-fragment complementation assays (PCAs). Plasmids encoding Gluc1-PIP4K2C and individual Gluc2-tagged viral proteins were transfected pairwise into HEK-293T cells followed by luciferase assays. PIP4K2C bound NSP3 and NSP6, whereas its co-expression with other viral proteins yielded background-level signals (Fig. 6e). A cutoff value of >2.2 SDs (corresponding to an NLR of >10) relative to a random reference set composed of 15 noninteracting human protein pairs^43^ was chosen as the threshold to define positive interactions.

These findings provide evidence that PIP4K2C is recruited by a viral protein to control the decrease in autophagic flux during SARS-CoV-2 infection.

We propose a model wherein PIP4K2C binds SARS-CoV-2 NSP6 and regulates impairment of autophagic flux by altering the phosphoinosite composition on autophagosomal or lysosomal membranes (Fig 6f). By acting on other membranes, PIP4K2C regulates various stages of the viral life cycle. RMC-113 induces PIP4K2C-mediated autophagy and PIKfyve-mediated viral fusion, thereby simultaneously targeting two key pathways implicated in SARS-CoV-2 and other viral infections.

## Discussion

There is an urgent need for novel antivirals to combat emerging viral infections and prepare for future pandemics. Impaired autophagic flux is one conserved cellular mechanism used by various viruses to support viral replication^20–23^, yet it is currently not targeted pharmacologically. Here we integrate virologic, pharmacological, genetic, biochemical, single cell transcriptomics, proteomics, and advanced lipidomics approaches with functional assays. We discover PIP4K2C as a key regulator of SARS-CoV-2 infection, interacting with NSP6, and uncover its role in regulating the virus-induced impairment of autophagic flux, providing insight into the underlying mechanism. Moreover, we validate PIP4K2C as a druggable antiviral target and reveal the therapeutic potential and mechanism of antiviral action of dual inhibition of PIP4K2C and PIKfyve as a novel broad-spectrum antiviral strategy.

We discover PIP4K2C, a lipid kinase not previously implicated in viral infections, as a candidate proviral factor using a pharmacological probe, RMC-113, whose targets we identify are PIP4K2C and PIKfyve based on unbiased kinome screens, targeted biochemical assays, pull-down by a click analog and docking/simulations. We validate a requirement for PIP4K2C in SARS-CoV-2 infection both genetically and via a chemically distinct selective PIP4K2C inhibitor. Utilizing a panel of assays with WT SARS-CoV-2, pseudovirus and subgenomic replicon, we reveal a requirement for PIP4K2C in viral entry, RNA replication and a late (assembly/egress) stage, likely facilitated by its presence on various membranes including plasma, autophagosomes, and Golgi^15^. Such control at different viral life cycle stages point to PIP4K2C as a key regulator of SARS-CoV-2 infection and an attractive antiviral target.

The three pandemic coronaviruses have been shown to impair autophagic flux, such as by blocking autophagosome–lysosome fusion and reducing lysosomal acidification^20, 21, 39, 44^, but the cellular mechanism that regulates this process remained unknown and untargeted. The reduced p62 and LC3-II protein degradation and autolysosome-to-autophagosome ratio we observed upon SARS-CoV-2 infection support virus-induced autophagic-flux impairment.

Extending previous findings showing alterations in autophagy gene expression^39^, the viscRNA-seq analysis on ALOs reveals temporal regulation of autophagy by SARS-CoV-2 with prominent activation of autophagy genes at an entry stage followed by prominent downregulation of autophagy and lysosomal genes at 24 hpi.

PIP4K2C is an understudied kinase, possibly due to its relatively low abundance and until recently, lack of pharmacological tools to probe its function. Prior reports demonstrating that its knockdown reduces autophagy cargo expression levels and mHTT aggregates in neurons and neuronal degeneration in HD Drosophila models indicate a role in autophagy^17, 18^. We provide pharmacological and genetic evidence that PIP4K2C is a key regulator of SARS-CoV-2-induced impairment of autophagic flux. Phenotypically, we show a partial reversion of the transcriptomic autophagy signature by viscRNA-seq upon RMC-113 treatment. Showing reduced p62 and LC3-II protein expression, along with an increased autolysosome-to-autophagosome ratio in SARS-CoV-2-infected cells upon PIP4K2C suppression via siRNA or RMC-113 (but not apilimod) treatment, functionally validates PIP4K2C’s role in autophagic flux impairment. It remains to be determined how suppression of autophagic flux by PIP4K2C promotes SARS-CoV-2 replication. Preventing degradation of viral proteins and/or increasing availability of autophagy factors for the formation of replication complexes^45, 46^ are among the possibilities.

The discovery that PIP4K2C binds SARS-CoV-2 NSP6 reveals its direct role in SARS-CoV-2 infection and suggests that it may regulate NSP6-mediated functions, such as lysosomal deacidification via ATP6AP1 binding^24^ and/or NSP6-mediated formation of non-digestive autophagosomes^46^. Intriguingly, probing the role of PIP4K2C in regulation of other interactions of SARS-CoV-2 proteins with autophagy factors shown to promote autophagosome-lysosome fusion impairment, such as ORF3a, ORF7a and NSP15^20, 47, 48^ is another interesting topic for future research.

Among the genes that are downregulated in SARS-CoV-2-infected cells relative to uninfected yet upregulated by RMC-113 treatment, particularly in VHCs, are the elongation factors *MAP1LC3B* (LC3B) and *SQSTM1 and WDFY3* whose encoded proteins (p62 and ALFY) form a complex tethered to autophagosome membranes, recruiting ubiquitinated protein aggregates^49^.

Contrastingly, the protein levels of p62 and LC3II increased following infection and decreased upon RMC-113 treatment. This indicates that while protein-level suppression of autophagic flus by SARS-CoV-2 is a primary result of infection, host cells might activate transcriptional downregulation to compensate for these virus-induced changes. In support of this hypothesis, cells within the infected culture that controlled infection effectively (zero vRNA counts) achieved greater suppression of SQSTM1 and WDFY3 expression than cells that failed to control infection (1+ viral counts).

The transcript levels of both *PLEKHM1* which facilitates autophagosome-lysosome fusion by interacting with components of the HOPS and SNARE complexes^50^, and *GABARAP*—a ubiquitin-like modifier on autophagosomes to which PLEKHM1 binds—were also downregulated in infected ALOs and upregulated upon RMC-113 treatment, yet with no correlation with vRNA presence. It is tempting to speculate that by reversing the autophagic flux impairment, RMC-113 promotes binding of ubiquitinated viral proteins, such as the nucleocapsid^51^, to these cargo receptors and their autophagic degradation. The observed upregulation of genes involved in various lysosomal functions upon RMC-113 treatment offers additional paths for future exploration of its mechanism of antiviral action.

Previous studies have tied phosphoinositide levels to autophagy regulation^52^: PI5P, PIP4K2C’s substrate, inhibits autophagy^18^. Moreover, viruses manipulate phosphoinositides to support their replication^13, 53, 54^. Using PRMC-MS lipidomics analysis, we comprehensively profiled various acyl chain variants of distinct PI classes, uncovering prominent alterations in phosphoinositide composition caused by SARS-CoV-2 infection. The PRMC-MS platform’s high-resolution revealed elevated levels of PI3P and PI5P--the lowest abundant and least studied phosphoinositide^55^—upon RMC-113 treatment, linking the antiviral effect to the suppression of PIKfyve and PIP4K2C and the modulation of autophagic flux^18^.

Beyond changes in PI3P and PI5P levels in infected cells, we provide additional evidence that modulating the enzymatic activity of PIP4K2C and PIKfyve is an important antiviral mechanism of RMC-113. RMC-113 mimics siPIP4K2C in suppressing SARS-CoV-2 entry, RNA replication, and infectious virus production, and in reversing the autophagic flux impairment. Moreover, WT PIKfyve and PIP4K2C, but not their kinase-dead mutants, reverse the anti-SARS-CoV-2 effect of RMC-113.

In contrast to PIP4K2C, PIKfyve’s role in various RNA viral infections and as a candidate antiviral target has been previously reported^12, 56^. We extend prior findings by showing antiviral activity of apilimod in ALOs and excluding a role for PIKfyve in stages beyond SARS-CoV-2 entry. While its role in endosome homeostasis is established, conflicting data exists regarding PIKfyve’s effect on autophagy regulation, with some studies suggesting induction^57, 58^ and others indicating suppression, at lower inhibitor concentrations^59^. This may explain our finding that apilimod, but not siPIKfyve, increased viral RNA replication and why apilimod and two other PIKfyve inhibitors could not clear the virus or protect mice from SARS-CoV-2 infection^60^.

Contrastingly, apilimod has demonstrated protection against influenza and RSV in mouse models^61^ and an excellent safety profile in human clinical trials for inflammatory diseases, de-risking PIKfyve as a target^9, 10^. Apilimod’s suboptimal pharmacokinetic profile^62^ has, however, limited its clinical development. Our data suggests that simultaneous suppression of PIKfyve and PIK4K2C by a single compound (i.e., “polypharmacology”) may improve effectiveness over suppression of PIKfyve alone. Nevertheless, while our data in ALOs are promising, the safety of PIP4K2C suppression remains to be determined.

By targeting two lipid kinases, RMC-113 inhibits replication of RNA viruses from four viral families (corona-, flavi-, alpha- and filoviruses). Moreover, passaging SARS-CoV-2 under RMC-113 treatment does not select for escape mutations, albeit the interpretation of these data in the absence of a DAA control (avoided to prevent gain of function mutations) is somewhat limited. Beyond viral infections, further development of RMC-113 and/or other PIP4K2C inhibitors may have implications for the treatment of non-infectious conditions in which PIP4K2C is implicated, including Huntington’s disease^17, 18, 63^.

In summary, our study discovers PIP4K2C, an understudied kinase, as a proviral factor required for SARS-CoV-2-mediated autophagic impairment and validates it as a druggable antiviral target. We propose dual inhibition of PIP4K2C and PIKfyve as a candidate therapeutic strategy for further development to enhance preparedness for viral outbreaks.

## Figure Legends

**Supplemental Fig. 1: RMC-113 has a broad-spectrum antiviral potential.**

**a,** Dose response to RMC-113 of rSARS-CoV-2-nLuc infection (black, USA-WA1/2020 strain, MOI = 0.05) and cell viability (blue) in Calu-3 cells via luciferase and alamarBlue assays at 24 hpi, respectively.

**b,** Dose response to RMC-113 of rVSV-SARS-CoV-2-S infection (black) and cell viability (blue) in Calu-3 cells via luciferase and alamarBlue assays at 24hpi, respectively.

**c and d,** Dose response of VEEV (TC-83) (MOI = 0.01) (**c**) and DENV2 (MOI = 0.1) (**d**) infections (black) and cell viability (blue) to RMC-113 measured in U-87 MG (**c**) and Huh7 (**d**) cells via luciferase and alamarBlue assays at 24 (**c**) and 48 (**d**) hpi, respectively.

**e and f,** Dose response of EBOV (Kikwit isolate, MOI = 1) (**e**) and MARV (Ci67 strain, MOI = 2)

(**f**) infections (black) and cell viability (blue) to RMC-113 measured in Huh7 cells via microneutralization and Cell-Titer-Glo luminescent assay at 48 hpi, respectively.

Data are a combination (**a-f**) or representative (**e, f**) of two independent experiments with 2-5 replicates each. Means ± SD are shown. Data are relative to DMSO control.

**Supplemental Fig. 2: RMC-113 is a selective kinase inhibitor that stably binds PIKfyve and PIP4K2C.**

**a,** Label-free kinase abundance ratio between RMC-113 (0.1, 1.0, and 10 µM)- and DMSO-treated SUM159 cell lysates measured following affinity purification via multiplexed inhibitor beads kinome profiling and analyzed by mass spectrometry (MIB/MS). Shown are the Log2 fold change mean values ± SD of 2 replicates.

**b,** Selectivity profiling of RMC-113 against 335 wildtype protein kinases at a concentration of 0.1 and 1 µM (ProQinase GmbH). Residual activity was determined relative to DMSO. The diameter of the dots reflects the % inhibition of the respective kinase. A selectivity score of 0.003 was measured. Data associated to ProQinase assays are shown in **Table S1**.

**c** and **d**, Positive electrospray ionization (ESI) mass spectra showing: (**c**) the protonated molecular ion peak of SRN2-002 at [M+2H]^+^ = 579.22 m/z, (**d**) a signally charged protonated molecular ion peak at [M+H]^+^ = 1194.56 m/z and the molecular ion peak at 1216.54 of the biotin-azide (1,2,3-triazole) clickable adduct. Insets depict the chemical structures and formulas of the detected peaks.

**e and f**, Induced fit docking pose of SRN2-002 in complex with (**e**) PIKfyve (PDB ID: 60SP), and (**f**) PIP4K2C (PDB ID: 2GK9). Residues within 3Å of the ligand are shown. Docking score (**e**) - 14.061 and (**f**) -11.836. Solvent exposed surface of the ligand is highlighted with yellow. H-bonds are shown with purple dashed lines; π–π interactions, green dashed lines, cation–π interactions, red dashed lines.

**Supplemental Fig. 3: Stabilization of MD simulations of RMC-113 in both PIKfyve and PIP4K2C.**

**a**, Root-mean-square fluctuation (RMSF) of RMC-113 heavy atoms in the simulation with PIKfyve (top) and PIP4K2C (bottom). The higher peak in *m*-MeO (atoms 7 and 8) illustrates the potential flip of the methoxy group in the binding site in the simulations.

**b**, Moving average RMSF 2D plot summarizing the observed protein–ligand interaction frequencies in the MD simulations. Interactions observed with >10% frequency are shown as bar plots. See 3D-locations of the residues in **Fig. 2**. H-bonds were defined as a 2.5Å distance with ≥120° angle for a donor and ≥90° for an acceptor, and water bridged interactions were defined as 2.8Å, ≥110° and ≥90°. The hydrophobic contact definition was 3.6Å for non-specific hydrophobic interactions and 4.5Å distance for ν–cation or ν–ν interactions. Simulation data of 72 μs (PIKfyve) and 20 μs (PIP4K2C) was analyzed for each 1 ns.

**c**, Root-mean-square deviation (RMSD) of PIKfyve and RMC-113 displayed for the concatenated simulation replica (18) trajectories. Simulation data of 72 μs was analyzed for each 1 ns, highlighting key interactions.

**d**, RMSD of PIP4K2C and RMC-113 displayed for the concatenated simulation replica (10) trajectories. Simulation data of 20 μs was analyzed for each 1 ns, highlighting key interactions.

**Supplemental Fig. 4: PIP4K2C and PIKfyve mediate the antiviral effect of RMC-113. a,** Schematic of the experimental procedures shown in **b**, **c**. ALO-derived monolayers were infected with WT SARS-CoV-2 (MOI = 1) and treated with apilimod.

**b,** Dose response to apilimod of SARS-CoV-2 infection (black) and cell viability (blue) in ALO supernatants via plaque and alamarBlue assays at 48 hpi, respectively.

**c,** Confocal IF microscopy images of F-actin (violet), SARS-CoV-2 nucleocapsid (green), and DAPI (blue) in SARS-CoV-2–infected ALOs pretreated with DMSO or apilimod (0.5 μM) at 24 hpi. Representative merged images at x40 magnification are shown. Scale bars: 50 μm.

**d** and **e,** Expression levels of PIKfyve (**d**), PIP4K2C (**e**), and actin via Western blotting at 24 hours post-transfection of Vero cells with control, WT PIKfyve (**d**) and PIP4K2C (**e**), or their kinase-dead mutant-expressing plasmids.

**f** and **g,** Vero cell viability measured by alamarBlue assays 48 hours post-transfection of the indicated plasmids.

**h,** Representative confocal microscopy images of A549-ACE2 cells transfected with mCherry-2xFYVE plasmid (red) 24 hours following treatment with DMSO, RMC-113 (5 μM) or apilimod (5 μM). Zoomed-in images at 40x magnification are shown. Scale bars are 10 μm.

**i** and **j,** Representative confocal microscopy images of A549-ACE2 cells transfected with mCherry-2xFYVE plasmid (red), treated with DMSO, RMC-113 (5 μM) or apilimod (5 μM) for 24 hours and stained for EEA-1 (green) (**i**) and Rab7 (green) (**j**). Representative merged and zoomed-in images at 40x magnification are shown. Scale bars are 10 μm.

Data represents a combination (**b,f**,**g**), one independent (**d, e, h**), or representative (**c, i, j**) of two independent experiments with 2-4 replicates each. Data in (**b, f, g)** are relative to DMSO. Means ± SD are shown.

**Supplementary Fig. 5: RMC-113 treatment alters the phosphoinositide regioisomer signatures.**

**a,** Levels of the indicated phosphoinositides (variant 38:4) as the fold change (FC) of peak area ratio in SARS-CoV-2-infected (MOI = 0.5) vs. uninfected Calu-3 cells, as measured via PRMC-MS.

**b-e,** Graphs are from the phosphoinositide lipidomics experiment shown in **Fig. 3 (j-m)**. FC in PI levels (all acyl chain variants except 38:4) for PI3P (**b**), PI5P (**c**) and their respective products PI(3,5)P_2_ (**d**), and PI(4,5)P_2_ (**e**) in SARS-CoV-2-infected cells are shown.

**f,** Levels of the indicated phosphoinositides (variant 38:4) expressed as the FC of peak area ratio in RMC-113- vs. DMSO-treated SARS-CoV-2-infected cells, as measured via PRMC-MS. **g**, Product-to-substrate ratio of the indicated phosphoinositidies in RMC-113- vs. DMSO-treated cells infected with SARS-CoV-2.

Data in **a, f, g,** are from an experiment that is independent of that shown in **Fig. 3j-m** and **Table S2.**

**Supplemental Fig. 6: Apilimod suppresses SARS-CoV-2 entry but increases RNA replication of a subgenomic SARS-CoV-2 replicon.**

**a,** Schematics of the experiments shown in **b** (**a**) and **d** (**c**).

**b,** Dose response to apilimod of rVSV-SARS-CoV-2-S infection (black) and cell viability (blue) in Calu-3 cells via luciferase and alamarBlue assays at 24 hpi, respectively.

**d,** Dose response to apilimod of viral RNA replication measured by luciferase assay in Vero E6-TMPRSS2 cells 24 hours post-transfection of in vitro transcribed nano-luciferase reporter-based SARS-CoV-2 subgenomic non-infectious replicon^38^.

Data is combination (**b, d**) of two independent experiments, each with 2–3 biological replicates. Data in **b, d** are relative to DMSO. Means ± SD are shown.

**Supplemental Fig. 7. RMC-113 promotes autophagic flux.**

**a**, Representative confocal microscopic images of uninfected A549-ACE2 cells transfected with GFP-RFP-LC3 tandem plasmid.

**b,** Abundance of autophagosomes (yellow) and autolysosomes (red) calculated as the total number of LC3 puncta per A549-ACE2 cell transiently transfected with GFP-RFP-LC3 tandem plasmid and infected with SARS-CoV-2 (MOI = 0.5) upon treatment with DMSO, RMC-113 (5 μM), apilimod (5 μM), or CQ (1 μM).

**c,** Representative confocal microscopy images of A549-ACE2 cells transfected with GFP-RFP-LC3 tandem plasmid and treated with DMSO, RMC-113 (5 μM), apilimod (5 μM), or CQ (1 μM), for 24 hours and stained for LAMP-1 (violet). Representative merged images at x40 magnification and zoomed-in images are shown. Scale bars: 10 μm.

Data is representative (**a,c**) or combination (**b**) of 2 independent experiments. Data in (**b**) is relative to DMSO. Means ± SD are shown. ****P <0.0001 by 1-way ANOVA followed by Dunnett’s multiple comparison test (**b**). ns= non-significant.

**Supplemental Fig. 8. RMC-113 differentially regulates autophagy-related genes in ALOs.**

**a,** Total number of cells in each condition at 24 hpi.

**b,** Heatmaps showing the log2 fold change in the expression of autophagy-related genes between: infected vs. uninfected (DMSO); RMC-113 vs. DMSO (uninfected); and RMC-113 vs. DMSO (infected) in AT2-like cells at 4 hpi. See **Fig.5l**. Black rectangles highlight transcripts with significant changes measured by Wilcoxon test.

**c** and **d,** Box plots showing the expression level of the indicated genes measured in AT2 cells at 24 hpi. Each dot represents an individual cell. Horizontal lines in each box indicate the median gene expression. P values by two-sided Wilcoxon test with Benjamini-Hochberg correction are shown.

**e** and **f,** Box plots depicting the differential expression of autophagy-related genes in AT2-like cells from SARS-CoV-2 infected ALOs at 24 hpi. Data are color-coded based on the median log2 change between VHCs and bystander cells in (**e**) DMSO-treated ALOs and (**f**) RMC-113 treated ALOs. Box plots: the horizontal lines indicate the first, second (median) and third quartiles; the whiskers extend to ±1.5× the interquartile range; the diamonds indicate outliers; the dotted vertical lines indicate ±1-fold change.

Data is a combination (**a-f**) of 3 independent experiments. *P* values by two-sided Wilcoxon test with Benjamini-Hochberg correction are shown.

**Supplemental Fig. 9. RMC-113 differentially regulates lysosomal genes in ALOs.** Heatmaps showing the log2 fold change in the expression of autophagy-related genes between infected vs. uninfected (DMSO) and RMC-113 vs. DMSO (uninfected) in AT2-like cells at 24 hpi. Black rectangles highlight transcripts with significant changes measured by Wilcoxon test.

**Supplemental Fig. 10. Suppression of PIP4K2C induces autophagic flux.**

**a,** Confirmation of siRNA-mediated knockdown by western blot in A549-ACE2 cells. Shown are protein expression levels at 48 hours post-transfection.

**b,** Representative confocal microscopic images of ACE-2-A549 cells transfected with the indicated siRNAs.

**c,** Abundance of autophagosomes (yellow) and autolysosomes (red) calculated as the total number of LC3 puncta per A549-ACE2 cell transfected with the indicated siRNAs.

Representative merged images at x40 magnification and zoomed-in images are shown. Scale bars: 10 μm.

Data is representative (**a, b**) or combination (**c**) of 2 independent experiments. Data in **c** are relative to DMSO. Means ± SD are shown. **P < 0.001, ****P <0.0001 by ordinary 1-way ANOVA followed by Dunnett’s multiple comparison test (**c**).

## Supplemental Table

**Table S1**. ProQinase data for RMC-113 at a concentration of 0.1 and 1 µM.

**Table S2.** Lipidomics data for two independent experiments.

**Table S3.** Modeling of PIKfyve and PIP4K2C.

## Materials and Methods

### Compounds

RMC-113 (Steven De Jong)^26^, Apilimod (Selleckchem, S6414), Rapamycin (Med chem express, #HY-10219), Bafilomycin A1 (Invivogen, #tlrl-baf1), CQ Chloroquine diphosphate (Premo Autophagy Tandem Sensor RFP-GFP-LC3B Kit, Thermo Scientific, # P36239). THZ-P1-2, CVM-05-002 and TM-04-176-01^33^ were a gift from Dr. Nathanael S. Gray, Stanford University.

### Plasmids

Plasmids used to produce SARS-CoV-2 pseudovirus were a gift from Jing Lin (Vitalant, San Francisco)^64^. The rSARS-CoV-2/WT and rSARS-CoV-2/Nluc (rSARS-CoV-2 expressing Nluc-reporter gene) plasmids were a gift from Dr. Luis Martinez-Sobrido ^65^. DENV2 (New Guinea C strain) TSV01 Renilla reporter plasmid (pACYC NGC FL) was a gift from Dr. Pei-Yong Shi (University of Texas Medical Branch)^66^. mCherry-2xFYVE plasmid was obtained from Addgene (#140050). SARS-CoV-2 subgenomic ΔS-E-M replicon was provided by Dr. Judith Gottwein (Copenhagen University Hospital, Denmark). VEEV-TC-83-nLuc RNA (a gift from Dr. William Klimstra (Department of Immunology, University of Pittsburgh, Pittsburgh). GFP-hPIKfyve (Addgene, #121148) cloned in gateway entry plasmid pDON221 and pDONR223-PIP4K2C (Addgene, #23450). Open reading frames encoding viral proteins (a gift from Dr. Peter Jackson, Stanford University) were recombined into a gateway-compatible pGluc fusion expressing vectors using Gateway technology (Invitrogen). Mutations were introduced by site-directed mutagenesis using the QuikChange Lightning Site-Directed Mutagenesis Kit (Agilent).

### Cells

Vero E6 cell line engineered to constitutively express enhanced green fluorescent protein (eGFP) was provided by Dr. Marnix Van Loock (Janssen Pharmaceutica, Beerse, Belgium) and was maintained in Dulbecco’s modified Eagle’s medium (DMEM, Gibco) supplemented with 10% v/v fetal calf serum (FCS, Biowest), 0.075% sodium bicarbonate and 1% penicillin-streptomycin (Pen-strep, Gibco). Vero E6, Vero, Calu-3, HEK-293T, U-87 MG, and BHK-21 cells (ATCC) were maintained in DMEM supplemented with 10% fetal bovine serum (FBS, Omega Scientific, Inc), 1% L-glutamine, 1% Pen-strep, 1% nonessential amino acids (NEAA, Gibco), 1% HEPES (Gibco), 1% Sodium pyruvate (Thermo Fisher Scientific). Vero E6-TMPRSS2 (JCRB cell bank, #JCRB1819) and A549-ACE2 cells (BEI resources, NR-53821) were maintained in DMEM supplemented with 10% FBS, 1% Pen-strep, and 1 mg/ml G418 (Gibco, #10131035). All cells were maintained in a humidified incubator with 5% CO_2_ at 37°C and tested negative for mycoplasma by MycoAlert (Lonza, Morristown, NJ).

### Human adult lung organoids (ALOs) model

The ALO model was generated from adult stem cells isolated from deep lung biopsy specimens. This model consists of cell types found in both proximal and distal airway epithelia, as validated previously^27^. Lung-organoid-derived monolayers were prepared as outlined^3, 27^ and plated in Pneumacult Ex-Plus Medium (StemCell Technologies).

### Cell viability assays

Cell viability was assessed using alamarBlue reagent (Invitrogen) according to the manufacturer’s protocol. Fluorescence was detected using GloMax Discover Microplate Reader (Promega).

### Viral stocks preparation and sequencing

The Belgium-GHB-03021 strain of SARS-CoV-2 was isolated from a nasopharyngeal swab obtained from a patient returning from China in early February 2020^67^ and passaged 6 times on Vero E6 cells. Viral stocks for rSARS-CoV-2/WT and rSARS-CoV-2/Nluc were generated as previously described^3^. Viruses produced in Vero E6-TMPRSS2 cells and passaged 3-4 times were used for the experiments. SARS-CoV-2 whole-genome sequencing was performed as previously described^3^, and showed no deletions in the spike multi-basic cleavage (MBC) domain. VEEV-TC-83-nLuc RNA was transcribed in vitro from cDNA plasmid templates linearized with MluI via MEGAscript SP6 kit (Invitrogen #AM1330) and electroporated into BHK-21 cells. DENV RNA was transcribed in vitro from pACYC-DENV2-NGC plasmid using mMessage/mMachine kits (Ambion) and electroporated into BHK-21 cells. EBOV (Kikwit isolate) and MARV (Ci67 strain) (BEI Resources) were grown in Vero E6 cells. Supernatants were collected, clarified, and stored at -80 °C, and viral titers were determined via plaque assays on BHK-21 (DENV, VEEV) or Vero E6 cells (SARS-CoV-2, EBOV, MARV).

For rVSV-SARS-CoV-2-S pseudovirus production, HEK-293T cells were transfected with spike expression plasmid followed by infection with VSV-G pseudotyped ΔG-luciferase VSV virus and harvesting of culture supernatant, as described^3^. rVSV-SARS-CoV-2-S was titrated via luciferase assay, and TCID_50_ was determined on Vero cells.

### Infection assays and pharmacological inhibition

Unless otherwise specified, inhibitors or DMSO were added 1-2 hours (cells) and 4 hours (ALOs) prior to viral inoculation and maintained throughout the experiment. Calu-3, Vero cells, or ALOs were infected with SARS-CoV-2 (MOI=0.05 or 1) in DMEM containing 2% FCS or 1X PneumaCult™-Ex Plus Medium at 37°C under biosafety level 3 (BSL3) conditions. Following a 1 to 4-hour incubation, the viral inoculum was removed, and cells were washed thoroughly and supplemented with fresh medium. At various time points postinfection, culture supernatants were collected for measurement of viral titer using standard plaque assays^68^, and cells were lysed in TrizolLS for RT-qPCR analysis. Huh7 cells were infected with DENV2 in (MOI=0.05) and at 48 hpi, viral replication was measured via luciferase assays. Huh7 cells were infected with EBOV (MOI=1) or MARV (MOI=2) under BSL4 conditions. At 48 hpi, cells were formalin-fixed for 24 hours prior to removal from BSL4 facility. Infected cells were detected using specific monoclonal antibodies against EBOV (KZ52) or MARV (7E6) glycoproteins and quantitated by automated fluorescence microscopy using an Operetta High Content Imaging System (PerkinElmer).

### Antibodies

*Immunoblotting:* Antibodies targeting PIKfyve (Thermo Scientific, #PA5-75977), PIP4K2C (Sigma, #WH0079637M1, Sigma), SQSTM1/p62, (Cell Signaling Technology, #5114), LC3A/B (D3U4C) XP® Rabbit mAb (Cell Signaling Technology, #12741), and β-actin (Sigma-Aldrich, #A3854).

*Immunofluorescence:* Antibodies targeting SARS-CoV-2 nucleocapsid (SinoBiological, #40143-MM05), EEA-1(Abcam, #ab109110), anti-LAMP1 (Abcam, #ab24170,), Rab7 (Origene, #AB0033-200). Goat anti-Mouse IgG (H+L), cross-absorbed secondary antibody, Alexa Fluor™ 633 (Thermo Scientific # A-21052A-21050), Goat anti-rabbit IgG (H+L), cross-absorbed secondary antibody, Alexa Fluor™ 488 (Thermo Scientific #A-11008).

### RNA interference

ON-TARGETPlus siRNA against PIKfyve, PIP4K2C, and SMARTpools non-targeting siRNA (siNT) (D-001206-13-05), were purchased from Dharmacon/ Horizon Discovery. siRNA sequences: PIKfyve – GGAAAUCUCCUGCUCGAAAUU; PIP4K2C - CCGAGUCAGUUGGACAACGAUU.

### siRNA transfection

siRNAs (10 pmol/well) were transfected into Calu-3, Vero E6 or A549-ACE2 cells using Dharmafect-4 (Dharmacon, #T-2004-02), lipofectamine RNAiMAX (Invitrogen) or Polyplus-transfection INTERFERin® (Genesee Scientific, #55-129), respectively, 48 hours prior to viral infection.

### Rescue assays

Plasmids encoding PIKfyve, PIP4K2C, or controls, were transfected into Vero cells using Lipofectamine 3000 reagent (Invitrogen) 24 hours before treatment and viral infection. Viral infection and cell viability were measured 24 hpi via luciferase and alamarBlue assays, respectively.

### RT-qPCR assays

Calu-3 cells were infected with SARS-CoV-2 (MOI = 1). At 2 hpi, the viral inoculum was removed, and cells were washed three times with PBS. Cells were then lysed in TRIzolLS (Invitrogen) and intracellular viral RNA levels were measured by RT-qPCR. The total RNA was extracted from cell lysates using Direct-zol RNA Miniprep Plus Kit (Zymo Research) and reverse-transcribed using High-Capacity cDNA RT kit (Applied Biosystems), following the manufacturer’s instructions. Primers and PowerUp SYBR Green Master Mix (Applied Biosystems) were added to the samples, and PCR reactions were performed with QuantStudio3 (Applied Biosystems). Target genes were normalized to GAPDH. Sequences of primers used for RT-qPCR are available upon request.

### Time-of-addition experiment

Calu-3 cells were infected with SARS-CoV-2 (MOI=1). Following 2 hpi for SARS-CoV-2), the virus inoculum was removed, and cells were washed twice with PBS. At specific time intervals, 10 μM RMC-113 or apilimod and 0.1% DMSO were added. Cell culture supernatants were collected at 10 hpi (SARS-CoV-2), and infectious viral titers were measured by plaque assays.

### Resistance studies

Vero E6-TMPRSS2 cells were infected with SARS-CoV-2 (MOI=0.05) and passaged daily 9 times by transferring an equal volume of viral supernatant to naive cells under DMSO or RMC-113 treatment at concentrations between the EC_50_ and EC_90_ values: 0.1 μM, passage 1; 0.3 μM passages 2-9. Viral titers in culture supernatants were measured by plaque assays.

### SARS-CoV-2 subgenomic replicon assay

The ΔS-E-M Nluc replicon was generated as described in^38^. Briefly, full-length DNA was linearized with NotI, followed by purification with the Zymo DNA clean & concentrator-25 kit but using Zymo-Spin™ IC-XL columns (ZR BAC DNA Miniprep Kit, ZymoResearch, Irvine, CA, USA). The linearized plasmid was in vitro transcribed using the mMESSAGE mMACHINE T7 Transcription Kit (ThermoFisher, Waltham, MA, USA) following the manufacturer’s instructions. RNA transcripts were quantified using the Qubit RNA BR Assay Kit (ThermoFisher, Waltham, MA, USA) and used for transfection. To measure RNA replication, in vitro transcribed replicon RNA was transfected into Vero E6 cells using Lipofectamine® RNAiMAX (ThermoFisher, #13778075), 48 hours prior to treatment with RMC-113 or following siRNA transfection. Viral RNA replication was measured after additional 24 hours via nano-luciferase assays.

### Immunoblotting

Whole-cell lysates were harvested in RIPA lysis buffer (Thermo Scientific) containing protease and phosphatase inhibitor cocktails. Protein concentration was measured using the detergent-compatible (BCA) protein assay (Thermo Scientific). Denatured lysates were separated on NuPage 8-16% Tris-glycine Midi Protein gels (Invitrogen) and transferred to polyvinylidene difluoride membrane (Immobilon) using a TransBlot Turbo dry transfer machine (Bio-Rad). The membrane was blocked and incubated with primary antibodies overnight at 4 °C followed by incubation with respective secondary antibodies and revealed with ECL Prime (Thermo). Chemiluminescent signals were acquired using Odyssey imaging system (Li-Cor), and densitometric analysis was performed using Image Studio 5.2 software (LI-COR, USA). All experiments were conducted and analyzed in 3-4 independent replicates.

### Immunofluorescence and confocal microscopy

For detection of viral protein in ALOs, SARS-CoV-2 infected ALO-derived monolayers were washed with PBS, fixed with 4% PFA, blocked, and incubated with mouse mAb SARS-CoV-2 nucleocapsid antibody (SinoBiological) overnight at 4 °C, followed by incubation with secondary antibodies, and counterstaining with DAPI (ThermoFisher). Images were taken on an SP8 microscope (Leica).

For colocalization and autophagy studies, A549-ACE2 cells were transduced with the Premo Autophagy Tandem Sensor RFP-GFP-LC3B kit (P36239, Thermo Scientific) and in 24 hours, infected with SARS-CoV-2 (MOI = 0.5) for 2 hours, followed by PBS washing and addition of medium containing 5 µM RMC-113, 5 µM apilimod, 1 µM chloroquine or DMSO. At 24 hpi, cells were fixed with 4% PFA, permeabilized with 0.5% (v/v) Triton-X-100, blocked with 2% (v/v) bovine serum albumin (BSA; Sigma-Aldrich) and stained as described above.

For imaging in siRNA-depleted cells, A549-ACE2 cells were transduced with premoRFP-GFP-LC3B sensor for 24 hours and infected with WT SARS-CoV-2 for 2 hours, washed with PBS, and replenished with the new medium. Images were acquired using LSM 710 confocal microscope (Zeiss).

To quantitate autophagic flux, image stacks were processed using 3D deconvolution in Zen 2.1 software. Maximum intensity projections of image stacks with both green and red channels were generated, and puncta counts for the respective channels were recorded using the Mosaic and ITCN plugins in Image J. Autolysosome l numbers were computed by subtracting the autophagosome count from the total puncta count, determined by combining the green and red channel image stacks using ImageJ.

### In vitro kinase assays

In vitro kinase assays to determine IC_50_ and dissociation constant (K_d_) were performed on the LabChip platform (Nanosyn) and Eurofins respectively.

**NanoBRET assays** were performed at Carna Biosciences. Briefly, HEK293 cells were transiently transfected with the NanoLuc® Fusion DNA and incubated at 37 °C. Twenty hours post-transfection, NanoBRET™ tracer reagent and the compounds were added to the cells and incubated at 37 °C for 2 hours. Nanoluciferase-based bioluminescence resonance energy transfer (BRET) was measured using NanoBRET™ Nano-Glo® Substrate on a GloMax^®^ Discover Multimode Microplate Reader (Promega).

### Kinome profiling

Multiplexed Inhibitor Bead (MIB) affinity chromatography/MS analysis was performed as previously described^28, 29^. SUM159 cell lysates were incubated with either DMSO or the indicated concentration of RMC-113 for 30 min on ice. Kinase fragments were then detected and analyzed by mass spectrometry. The abundance of kinases was quantified in a label-free manner using MaxQuant software.

**ProQinase assay** was conducted by ProQinase (GmbH, Germany) by measuring residual activity of 335 wild-type protein kinases upon incubation with RMC-113 at 0.1 and 1 μM.

### Molecular modeling

Molecular modeling was conducted with Maestro (Schrödinger Releases 2021-4/2022-2, Maestro, Schrödinger, LLC, New York, NY, 2021/2022) with OPLS4 force field^69^. Induced fit docking of SRN2-002 to MD simulation derived structures was conducted as reported earlier^70^. MD simulations of RMC-113 in complex with PIKfyve (PDB ID: 7K2V)^71^ and PIP4K2C (8BQ4)^72^ were run with Desmond^73^, resulting in total aggregate simulation data of 72 μs and 20 μs, respectively.

### Molecular dynamics simulations

To build the PIKfyve–RMC-113 complex, as no high-quality crystal structure of PIKfyve kinase domain was available, we used the cryo-EM structure (PDB ID: 7K2V)^71^. RMC-113 was manually placed on the binding site of PIKfyve utilizing the information of an analogue structure in complex with the lipid kinase PI3K p110δ (PDB ID: 2WXM)^74^ (Schrödinger Release 2021-4). Next, the obtained protein–ligand complex structure was prepared with Protein Preparation Wizard^75^. The termini were capped, H-bonds were optimized, and system was energy minimized twice using 0.5Å heavy atom RMSD convergence. The energy minimized system was solvated in a cubic box with a 15Å minimum distance to the box edges from the protein or ligand atoms. We used TIP3P model^76^ to describe the water and system was neutralized with six Cl^-^ ions and K^+^ and Cl^-^ ions were added to obtain a final salt concentration of 0.15M. The final system contained a total of 55,180 atoms. The default Desmond relaxation protocol was applied before the production simulations. The production simulations were run in NpT ensemble: 1.01325 bar, Nosé–Hoover method; 300 K, Martyna–Tobias–Klein method; RESPA integrator with 2, 2, and 6 fs timesteps for bonded, near and far, respectively; Coulombic cutoff of 9Å. In total, we ran 20 replica simulations **(Table S3)**, each with a different random seed, with a length of 4 μs each. Within these final simulations, we discarded two replicas from further analysis due to their conformational instability. The remaining 18 replicas, with an aggregate simulation data of 72 μs, were used in the final analysis.

Building the PIP4K2C–RMC-113 complex, used a combination of x-ray structure 8BQ4^72^ and AlphaFold model (AF-Q8TBX8-F1)^77^ using (Schrödinger Release 2022-2). RMC-113 was superimposed over the co-crystallized ligand of 8BQ4 and the structure was directed to protein-ligand complex refinement protocol for local optimization of side chain conformations. A similar pose as to PIKfyve was gained. Missing loops of the template protein (residues Gly136–Gly139; Ile291–Phe350; Thr377–Thr402) were modelled with the assistance of Maestro’s homology modelling tool, using multiple template approach i.e., consensus protocol, where chains A and B from 8BQ4 and AlphaFold^9^ model AF-Q8TBX8-F1 were selected as input structures, respectively. As a result, a complete PIP4K2C kinase domain (residues 45-421) structure was obtained, where the core of the protein was taken from 8BQ4, and missing loops were obtained from AF-Q8TBX8-F1. The earlier obtained RMC-113 pose was merged to new model and the refinement step was repeated, after which the structure was H-bond optimized and minimized using standard Protein Preparation Wizard (as above). The energy minimized structure, was solvated with TIP3P waters in an orthorhombic periodic system with 10Å buffer (91.0 x 77.9 x 71.2 Å), and neutralized, including 0.1 M NaCl buffer. The final system contained a total of 46,285 atoms. Desmond MD simulations were carried out using same settings as with PIKfyve, resulting in 10 replicas of with the length of 2 μs each (total simulation time of 20 μs) **(Table S3).**

Simulation trajectories were analyzed by Maestro simulation interactions diagram tools and visualization of the structures was conducted with PyMOL (The PyMOL Molecular Graphics System, Version 2.5.4 Schrödinger, LLC.)

### Target engagement by Click chemistry

As described^31^, cell lysates were prepared from A549-ACE-2 cells infected with SARS-CoV-2 at 24 hpi (MOI = 1). Lysis buffer containing 50 mM PIPES, 50 mM NaCl, 5 mM MgCl2, 5 mM EDTA, 0.5% NP-40, 0.1% Triton X-100, and 0.1% Tween 20 at pH 7.4 was used to inactivate the virus, followed by centrifugation to remove debris. For the biotin-azide-streptavidin magnetic pull-down, cell lysates were incubated with the photoaffinity clickable probe (SRN2-002, 1 µM and 5 µM), followed by UV light irradiation. A CuAAC click reaction mixture containing CuSO4, THPTA, Biotin-Azide, and sodium ascorbate was added, and after desalting, streptavidin magnetic beads were used to pull down biotin-protein complexes. The resulting product was washed and analyzed by immunoblotting. Controls included samples without the clickable probe, without UV irradiation, and with competitive click reactions using excess parent compound (RMC-113, 10X or 50X).

### Lipidomics analysis. Lipid extraction, derivatization, and methylation

A549-ACE2 cells infected with SARS-CoV-2 (MOI=0.5) were treated with 5 µM RMC-113 or DMSO. At 24 hpi, cells were lysed, and samples were inactivated using 1.5 ml methanol and mixed with chloroform (chloroform/methanol 1:9), supplemented with 1 nmol of PI(4,5)P2 as an absorption inhibitor to prevent non-specific binding, along with synthetic phosphoinositides [PI(3,4,5)P3, PI(3,4)P2, PI(3,5)P2, PI(4,5)P2, PI3P, PI4P, PI5P, and PI] (10 pmol each) as internal standards. The mixture was inactivated with 1.5 ml methanol and chloroform, followed by addition of Ultrapure water, 2 M HCl, and 1 M NaCl. The crude lipid extract (2.9 ml) was subjected to purification using a DEAE Sepharose Fast Flow column, washed with chloroform/methanol (1:1) and chloroform/methanol/28% aqueous ammonia/glacial acetic acid (200:100:3:0.9), and then eluted with chloroform/methanol/12 M hydrochloric acid/ultrapure water (12:12:11). The eluate was combined with 850 µl of 120 mM NaCl and purified by centrifugation. The resulting purified phosphoinositides underwent derivatization through methylation by adding 0.6 M trimethylsilyl diazomethane at room temperature for 10 minutes. The reaction was quenched with 20 µl of glacial acetic acid. The samples were further processed with methanol/ultrapure water/chloroform (48:47:3), followed by evaporation under nitrogen and dissolution in 100 µl of acetonitrile for analysis.

### MS analysis using Chiral column chromatography and tandem mass spectrometry

These experiments were conducted at Element Materials Technology. The lipid samples were analyzed using chiral column chromatography coupled with tandem mass spectrometry (PRMC-MS), employing a QTRAP6500 mass spectrometer operating in positive ion mode. Mass spectra were acquired over a broad m/z range (5–2000), with multiple reaction monitoring (MRM) applied for targeted data acquisition. Instrument parameters, including ionization voltages (5.5 kV), gas pressures (20 psi for curtain gas, 9 psi for collision gas, and 50 psi for ion source gases), and temperature (100 °C source block temperature), were meticulously optimized to maximize sensitivity and resolution. Liquid chromatography (LC) separation was achieved using a Lux 3 µm i-Cellulose-5 column (250 x 4.6 mm) maintained at 35 °C, with a mobile phase gradient of methanol/5 mM ammonium acetate (A) and acetonitrile/5 mM ammonium acetate (B). The LC gradient started at 0% A and 100% B, transitioning to 30% A and 70% B over 3 minutes, held for 15 minutes, and then re-equilibrated. Data processing involved normalization against deuterated reference standards, enabling precise quantification and comparison of lipid species across samples.

### ViscRNA-seq

ALOs were seeded in 96-well plates at 1x10^5^ cells per well and infected with SARS-CoV-2 (MOI=1) in the presence of 5 μM RMC-113 or 0.1% DMSO. After a 4-hour incubation, the viral inoculum was removed, and cells were washed thoroughly and supplemented with fresh medium with or without the compound. At 4 and 24 hpi, ALOs were washed and then detached via 5-min incubation with TrypLE followed by centrifugation at 300 g for 10 minutes. The pellets were suspended and fixed with Cell fixation solution (Parse, Evercode Fixation, cat: ECF2001, Part number WF303) for 10 minutes followed by incubation with Cell permeabilization Solution (Parse, Evercode Fixation, cat: ECF2001, Part number WF305) for 3 minutes at RT. ALOs were then taken out from BSL3 for single-cell sequencing according to the manufacturer’s instructions (Parse, EvercodeTM WT mini Mega v2).

### viscRNA-seq data analysis

#### Data processing

The Parse Biosciences processing pipeline (split-pipe v1.0.3a) was used with default settings to align the raw sequencing reads to the Parse Biosciences pre-built human reference (GRCh38) and to demultiplex samples. The downstream processing was performed using the Python package Scanpy^78^. Cells with more than 30% mitochondrial reads or more than 50,000 unique molecular identifiers (UMIs) were excluded from subsequent analysis. The raw gene counts matrix was normalized to a total UMI counts of 1,000,000 (CPM) per cell and was log2 transformed with the addition of a pseudocount of 1.

#### Cell clustering and annotation

Samples from three independent experiments were clustered and annotated separately. Principal component analysis (PCA) was performed on the 2,000 features that were highly variable across most of the samples. The first 50 PCA components were used to generate the nearest neighbor graphs and were subjected to a non-linear dimensionality reduction using Uniform Manifold Approximation and Projection (UMAP). Cells were clustered using the Leiden algorithm with a resolution of 1. Cell clusters were annotated based on the expression of marker genes as follows: (1) AT2-like cells (SFTPD, MUC1, CLIC5), (2) AT1-like cells (HOPX, RTKN2, EMP2, CAV1), (3) Basal-like cells (NGFR, ITGA6, TP63, KRT5), and (4) transitioning cells that are HOPX and NGFR negative and CEACAM6 positive. After initial annotation, PCA components were computed on the 2,000 features that were highly expressed across all the samples from independent experiments. Principal components were adjusted for each sample using Harmony integrate function. The first 40 Harmony components were used to generate batch-corrected UMAP visualizations.

#### Detection of viral RNA harboring cells (VHCs)

To capture the viral RNAs in SARS-CoV-2 infected samples, the raw sequencing reads were aligned to a custom reference genome by integrating the SARS-CoV-2 isolate USA-WA1/2020 (MT246667.1) to the human reference genome (GRCh38) using the Parse Biosciences processing pipeline as described above. Reads uniquely mapped to the viral genome were extracted as viral reads. Cells with at least four detected viral transcripts were considered as VHCs, and cells with zero detected viral transcripts were considered as bystanders. This threshold was defined empirically based on the inflection point in the cumulative distribution of viral RNA counts across cells in each sample.

#### Pairwise comparison of VHCs and bystanders

Both VHCs and bystanders were subsampled randomly to the lowest number of cells in one group and log2 fold change of the gene expression was calculated. This calculation was iterated 100 times to form an empirical distribution of differential expression. The median log2 fold change of all the comparisons was used to estimate effect size (color) and the noise level (box size) of the comparison was shown using boxplots.

#### Protein-fragment complementation assays

As described, combinations of plasmids encoding prey (A) and bait (B) proteins, each fused to a fragment of the Gaussia luciferase protein (GLuc1 and GLuc2) or control vectors, were co-transfected into HEK-293T cells plated in 96-well plates in triplicate^43^. At 24 hours post-transfection, cells were lysed and subjected to luciferase assays (Promega). Results were expressed as NLRs calculated as follows: the average signal in cells transfected with GLuc1-A and GLuc2-B was divided by the average signal in wells transfected with GLuc1-A and an empty GLuc2 vector and those transfected with GLuc2-B and an empty GLuc1 vector.

#### Statistical analysis

Data were analyzed with GraphPad Prism software. EC_50_ and CC_50_ values were measured by fitting of data to a 3-parameter logistic curve. P values were calculated by 1-way ANOVA with either Dunnett’s or Tukey’s multiple comparisons tests or a two-sided Wilcoxon test with Benjamini-Hochberg correction as specified in each figure legend.

#### Study Approval

SARS-CoV-2 and filovirus work were conducted in BSL3 and BSL4 facilities at Stanford University, KU Leuven Rega Institute, and USAMRIID according to CDC and institutional guidelines. Human lung organoid (ALO) propagation was approved under protocol IRB# 190105 at UCSD.

#### Data availability

PDB coordinates of the representative snapshots shown in **Fig. 2** are provided as supplementary information files. Full trajectories of the Desmond molecular dynamics simulations are freely available at: https://doi.org/10.5281/zenodo.10878041 (PIKfyve simulations) and https://doi.org/10.5281/zenodo.10889319 (PIP4K2C simulations).

## Acknowledgements

This work was supported by grant 1R01AI158569-01 from the NIH (to SE), awards HDTRA11810039 from the Defense Threat Reduction Agency/Fundamental Research to Counter Weapons of Mass Destruction (to SE, SDJ) and W81XWH2210283 and W81XWH-16-1-0691 from the Department of Defense/Congressionally Directed Medical Research Programs (to SE and SDJ), and grants 3R01DK107585-05S1 and UCOP-R00RG2642 (to SD). SE is a Chan Zuckerberg Biohub investigator. This work was partly supported by the NIH Common Fund Illuminating the Druggable Genome (IDG) program (NIH Grant U24DK116204) (CRMA, GLJ, ME). MK was supported by a PhRMA Foundation Postdoctoral Fellowship in Translational Medicine and NIH T32 training grant (5T32AI007502-27). MM was supported by the Stanford Pandemic Preparedness Hub Fellowship. LG was supported by a long-term European Molecular Biology Organization Fellowship (ALTF584-2021).

Microscopy and mass-spectrometry were done on instruments in the Cell Sciences Imaging Core Facility (RRID:SCR_017787, LSM780 confocal/multiphoton) and Thermo Exactive Orbitrap LC/MS system (RRID:SCR_018700), respectively. We thank the investigators who provided plasmids and compounds (see Methods). We thank the Stanford BSL3 service center and Jaishee Garhyan (director) for their assistance in the BSL3. We thank Biocenter Finland/DDCB for financial support and CSC – IT Center for Science, Finland, for computational resources. We thank the Stanford Clinical Virology Laboratory’s staff for their help with sequencing. Opinions, conclusions, interpretations, and recommendations are those of the authors and are not necessarily endorsed by the funders. The mention of trade names or commercial products does not constitute endorsement or recommendation for use by the Department of the Army or the Department of Defense.

