## Supplemental Fig for "PIP4K2C inhibition reverses autophagic flux impairment induced by SARS-CoV-2"

### Supplementary Fig. 1

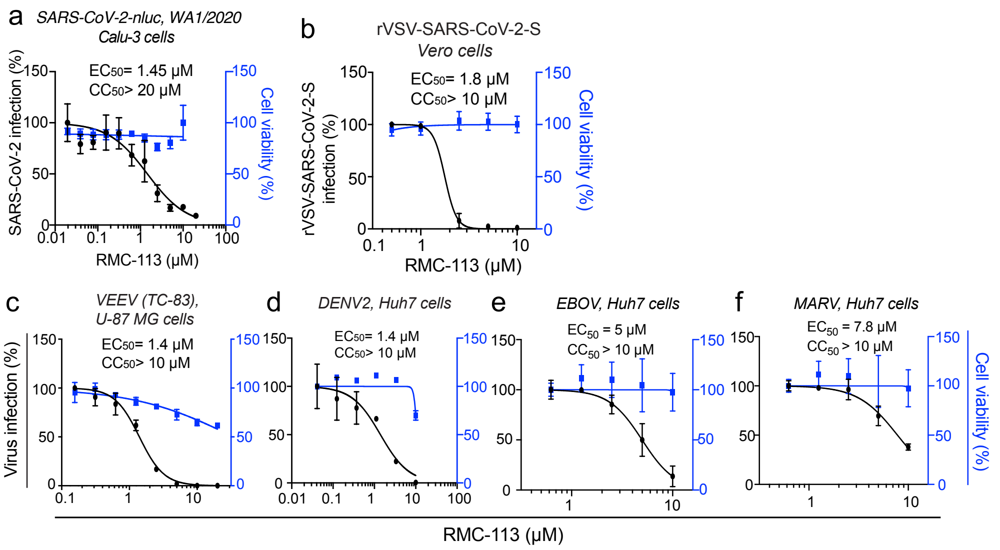

Supplementary Fig. 2

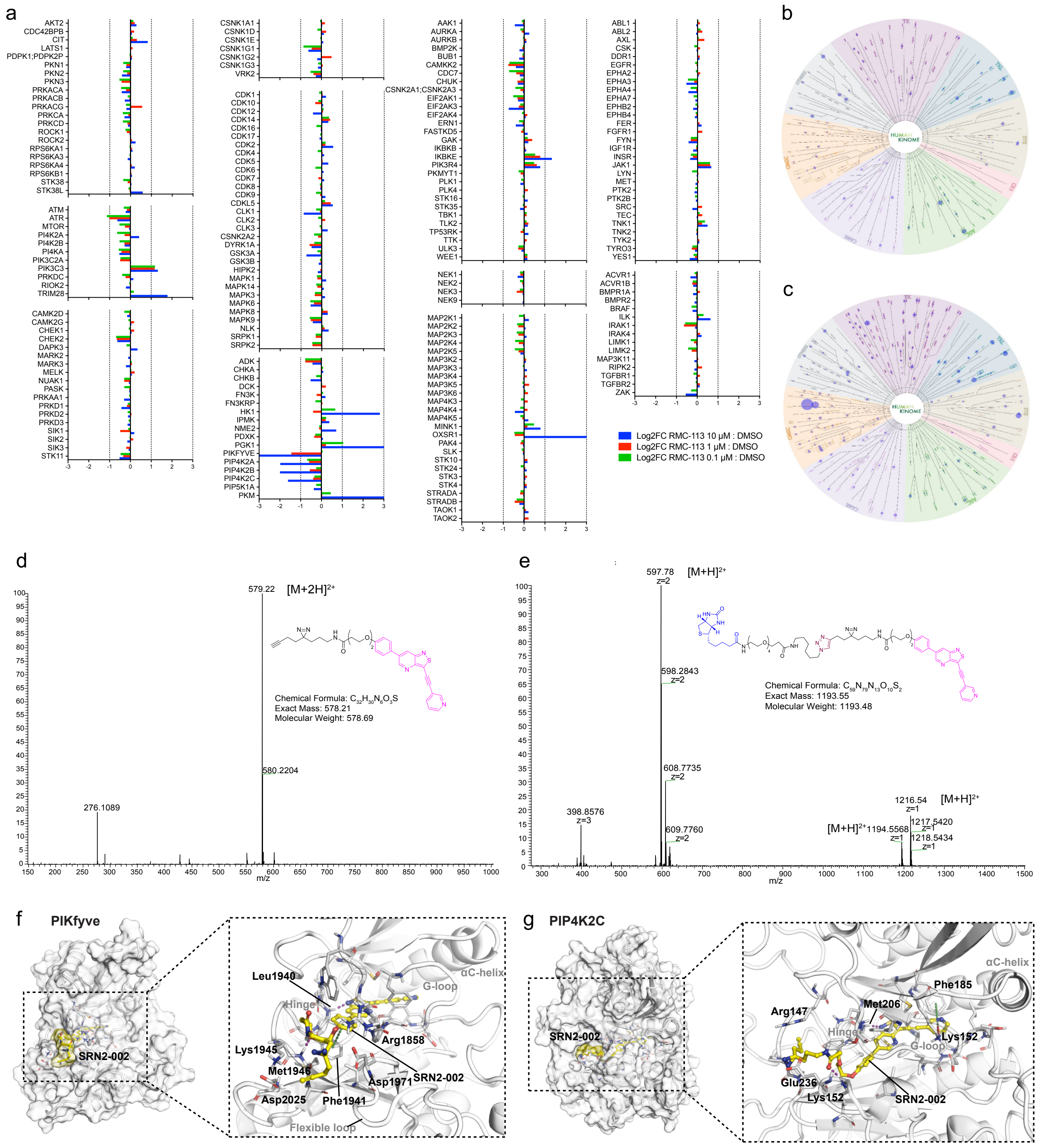

### Supplementary Fig. 3

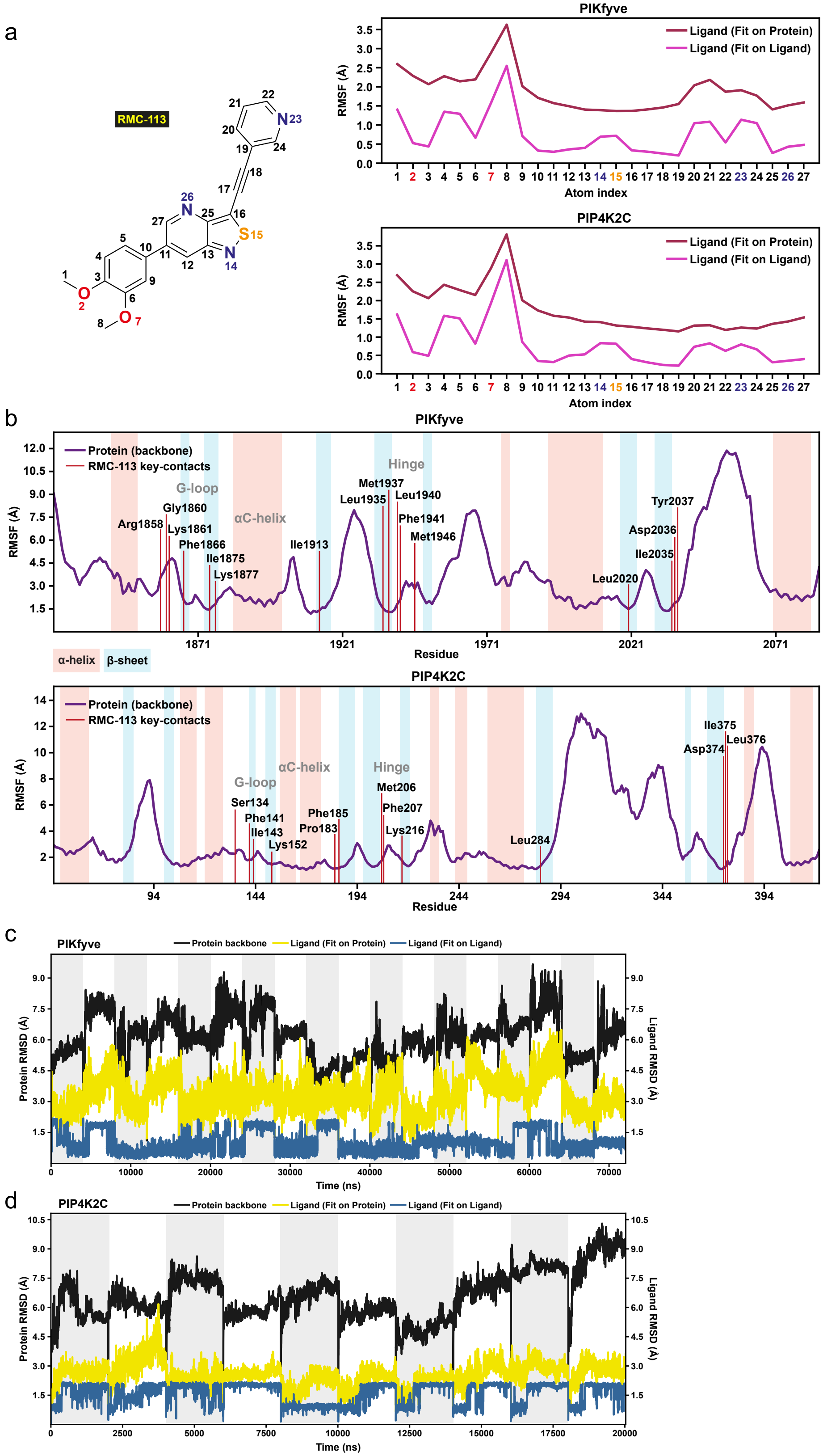

Supplementary Fig. 4

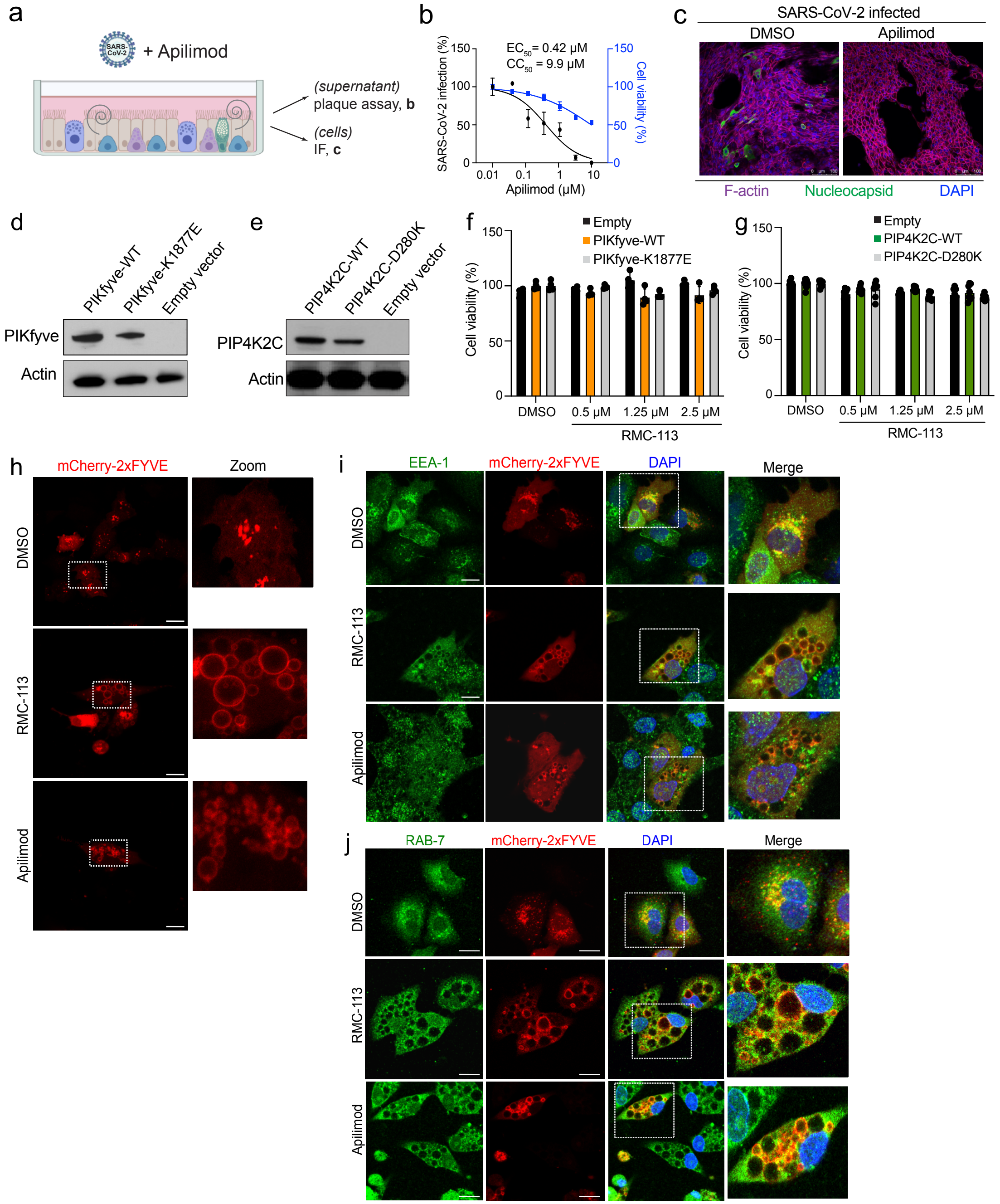

### Supplementary Fig. 5

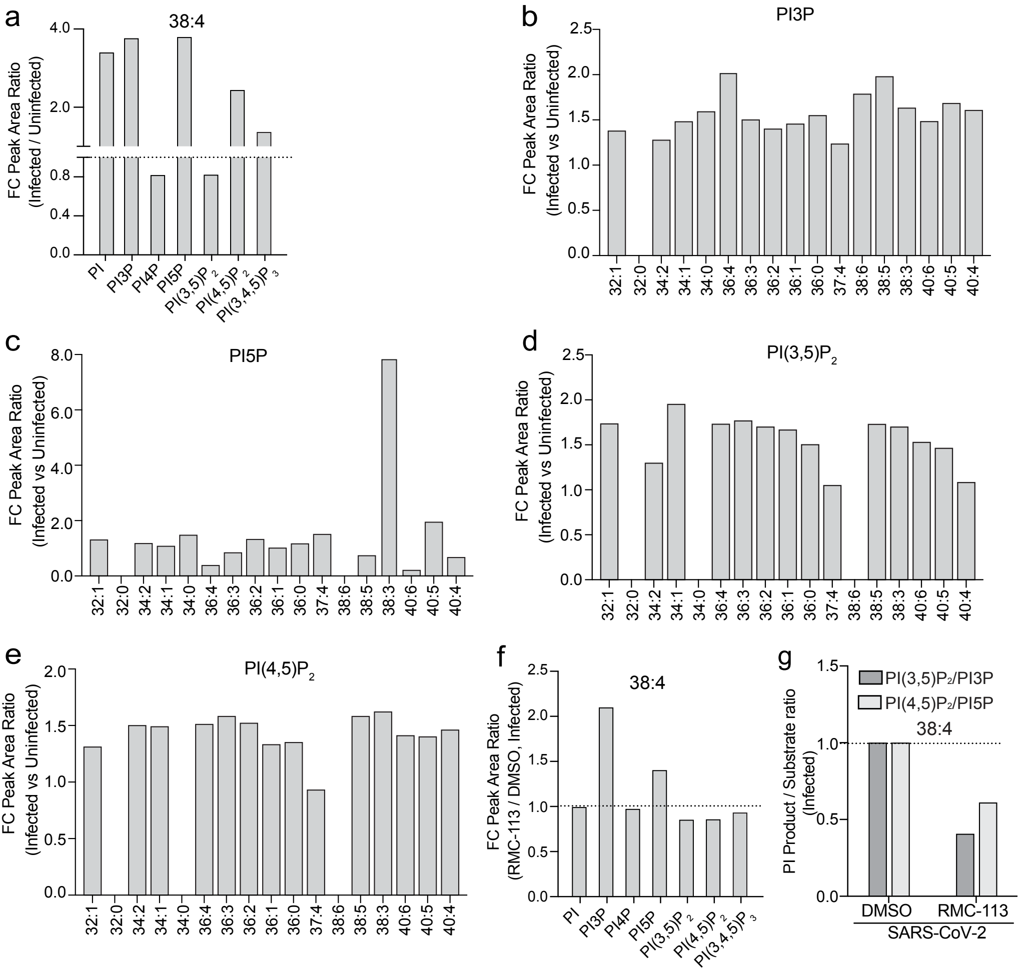

### Supplementary Fig. 6

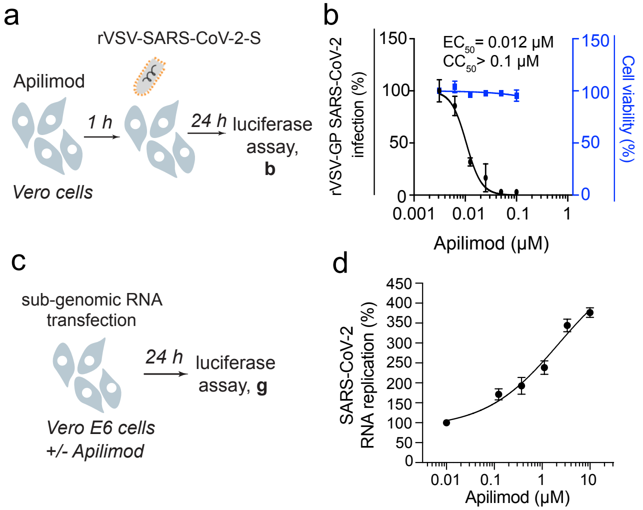

### Supplementary Fig. 7

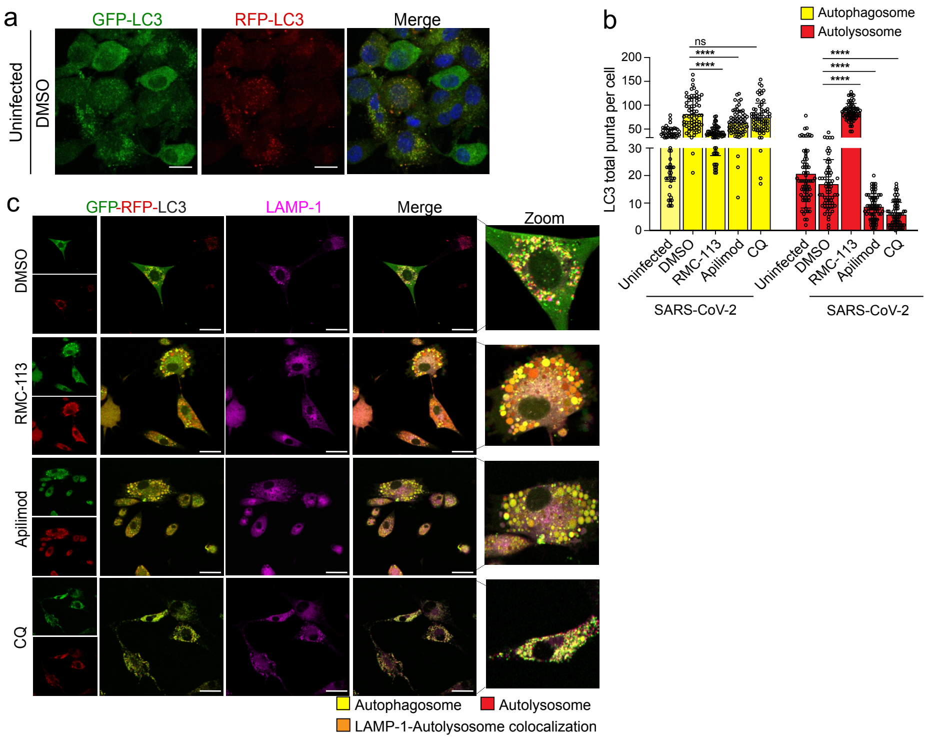

### Supplementary Fig. 8

**a**

| Infection status | Treatment | VHCs | | Bystanders |
|---|---|---|---|---|
| | | high >3 | low (1-3) | zero |
| SARS-CoV-2 | DMSO | 421 | 2649 | 5571 |
| | RMC-113 | 115 | 1363 | 6388 |
| Uninfected | DMSO | 0 | 0 | 1369 |
| | RMC-113 | 0 | 0 | 1833 |

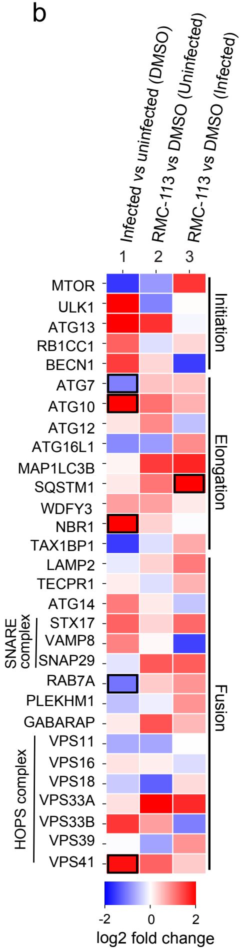


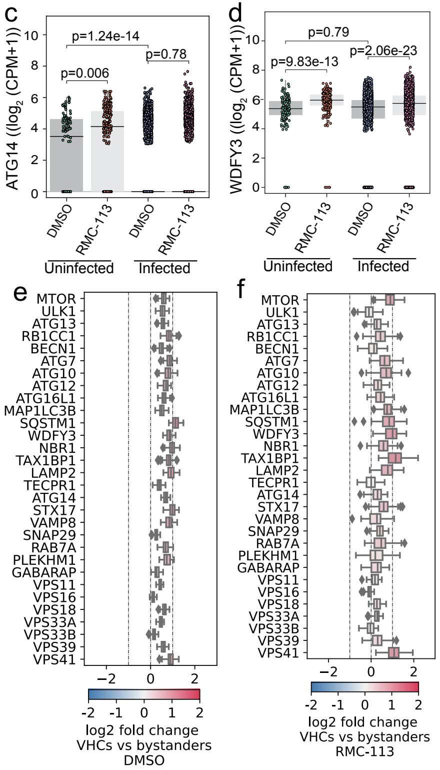

### Supplementary Fig. 9

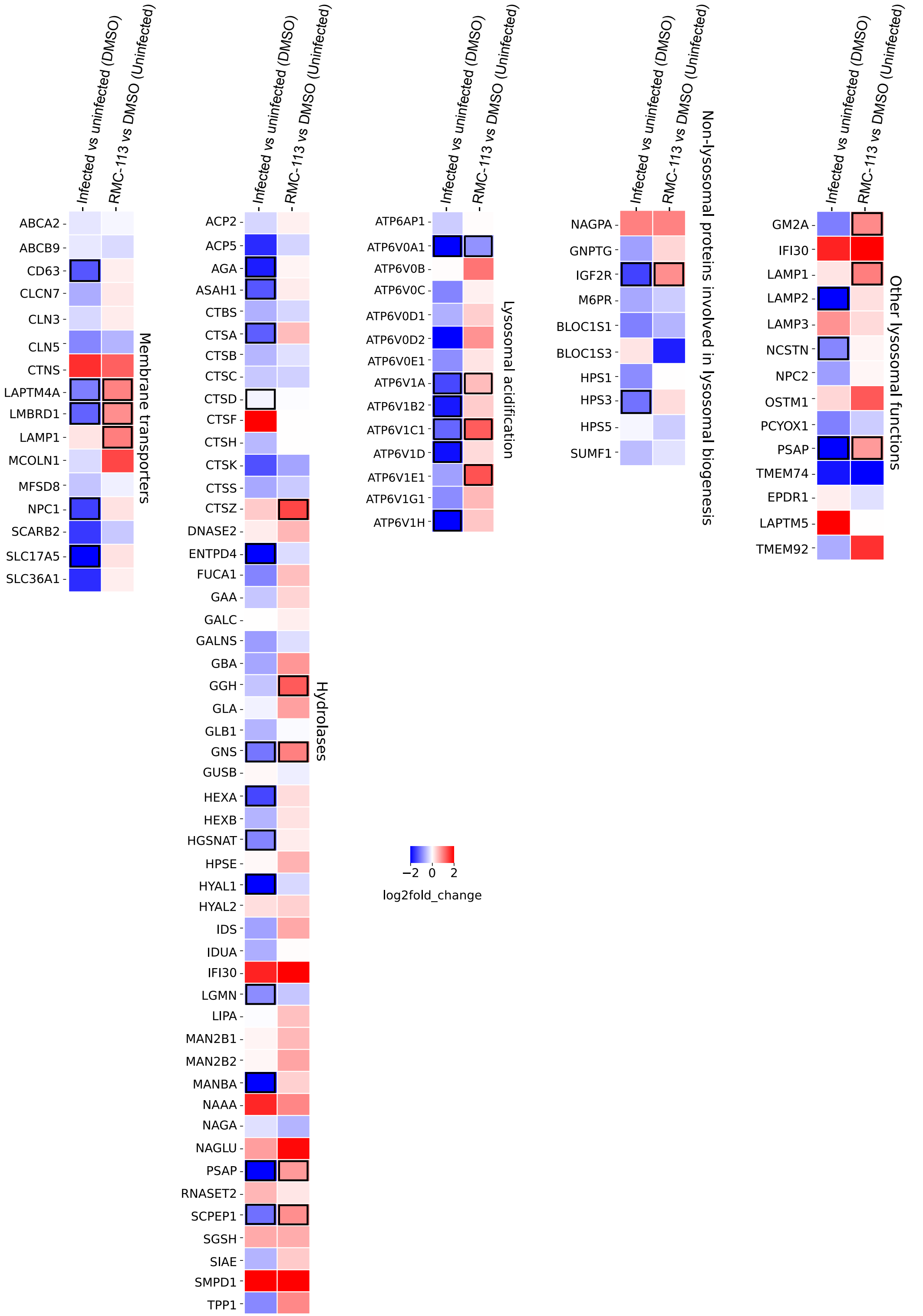

### Supplementary Fig. 10

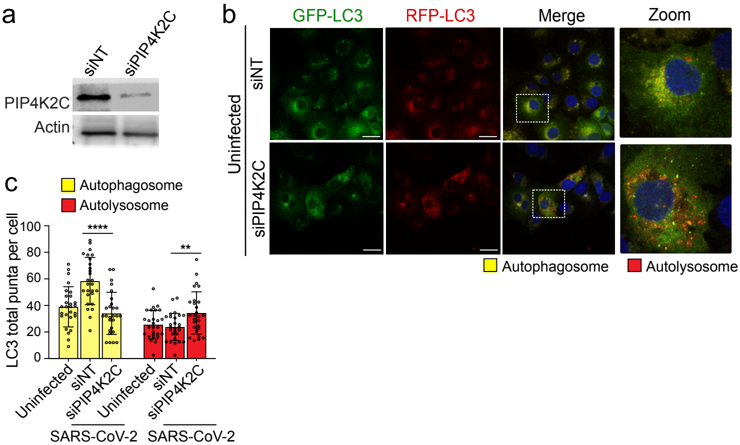
